# IDH1 Mutations Induce Organelle Defects Via Dysregulated Phospholipids

**DOI:** 10.1101/2020.03.20.000414

**Authors:** Adrian Lita, Artem Pliss, Andrey Kuzmin, Tomohiro Yamasaki, Lumin Zhang, Tyrone Dowdy, Christina Burks, Natalia de Val, Orieta Celiku, Victor Ruiz-Rodado, Elena-Raluca Nicoli, Michael Kruhlak, Thorkell Andresson, Sudipto Das, Chunzhang Yang, Rebecca Schmitt, Christel Herold-Mende, Mark R. Gilbert, Paras N. Prasad, Mioara Larion

## Abstract

Cytosolic IDH1 enzyme plays a key, but currently unexplored, role in lipid biosynthesis. Using Raman imaging microscopy, we identified heterogeneous lipid profiles in cellular organelles attributed uniquely to IDH1 mutations. Via organelle lipidomics, we found an increase in saturated and monounsaturated fatty acids in the endoplasmic reticulum of IDH1^*mut*^ cells compared with IDH^*WT*^ glioma. We showed that these fatty acids incorporate into phospholipids and induce organelle dysfunctions, with prominent dilation of Golgi apparatus, which can be restored by transient knockdown of stearyl-CoA desaturase or inhibition of D-2-hydroxyglutarate (D-2HG) formation. We validated these findings using tissue from patients with glioma. Oleic acid addition led to increased sensitivity to apoptosis of IDH1^*mut*^ cells compared with IDH^*WT*^. Addition of D-2HG to U251^*WT*^ cells lead in increased ER and Golgi apparatus dilation. Collectively, these studies provide clinically relevant insights into the functional link between IDH1^*mut*^-induced lipid alterations and organelle dysfunction, with therapeutic implications.

**Significance:** Gliomas are devastating tumors, with the most aggressive form—glioblastoma multiforme— correlated with a mean patient survival of 14.5 months. No curative treatment exists to date. Low-grade glioma (LGG) with the isocitrate dehydrogenase 1 (IDH1) mutation, R132H, provides a survival benefit to patients. Understanding the unique metabolic profile of IDH1^*mut*^ could provide clues regarding its association with longer survival and information about therapeutic targets. Herein, we identified lipid imbalances in organelles, generated by IDH^*mut*^ in cells and patient tissue, that were responsible for Golgi dilation and that correlated with increased survival. Addition of oleic acid, which tilted the balance towards elevated levels of monounsaturated fatty acids produced IDH1^*mut*^-specific cellular apoptosis.

**Highlights:**

- Single-organelle omics revealed unique alterations in lipid metabolism due to IDH1-mutations.
- IDH mutation leads to organelle-wide structural defects.
- IDH1 mutation leads to increased monounsaturated fatty acids levels in glioma cells and oligodendroglioma patient samples.
- Lipid alterations affect the membrane integrity of the Golgi apparatus.
- Increased D-2HG induced SCD expression and elevated monounsaturated fatty acids
- Tilting the balance toward more-abundant monounsaturated fatty acids leads to specific IDH1^*mut*^ glioma apoptosis.

## Introduction

Mutations of the isocitrate dehydrogenase 1 (IDH1) gene play an intriguing role in the development of glioma and other tumors (Waitkus et al., 2015, Parsons et al., 2008, Khan et al., 2017, Yan et al., 2009, Medeiros et al., 2017, Wang et al., 2018, Brunner et al., 2019, Mohammad et al., 2019, Lopez et al., 2010, Victor et al., 2019). IDH1 mutations are an early event, (Lass et al., 2012) are associated with a less aggressive phenotype (Parsons et al., 2008) potentially due to their slow growth and need for nutrients to form D-2-hydroxyglutarate (D-2HG), and are used as prognostic and diagnostic markers of glioma. In fact, the World Health Organization (WHO) released a novel classification of glioma in 2016 to include IDH1 mutations as molecular markers that dictate the classification (Louis et al., 2016). Much effort has been directed toward inhibiting D-2HG formation(Yen et al., 2010, Han and Batchelor, 2017); however, the links between IDH1 mutations, tumor metabolism, and clinical manifestation are not well understood.

Cytosolic NADP-dependent IDH1 plays an important role in lipid biosynthesis via its the production of citrate and NADPH (Koh et al., 2004). The loss of the wildtype (WT) allele in gliomas with the arginine 132 to histidine mutation leads to impaired citrate formation; moreover, the neomorphic activity of mutant IDH1 utilizes NADPH to synthesize up to 10 mM of D-2HG, which is a biomarker for tumor cells carrying an IDH1 mutation. The combined effect of loss of wild type allele and usage of NADPH by the mutated allele leads to relative depletion of those precursors of lipid biosynthesis. Despite the direct impact of IDH1 mutation on lipid biosynthesis, little is known about how altered lipid metabolism affects specifically IDH1^*mut*^ gliomas. Thus, we sought to determine the lipid profile changes due to IDH1 mutation at organellar levels utilizing our newly developed methodology.

Subcellular compartmentalization of metabolic processes is a major regulator of the overall cellular metabolome. Processes such as autophagy, for example, rely on lysosomal sensors (Wyant et al., 2017) to export essential amino acids out of the lysosome; oxidative phosphorylation of glucose takes place in the mitochondria; and the Golgi apparatus (GA) and endoplasmic reticulum (ER) play central roles in lipid metabolism (Bankaitis et al., 2012). Classical metabolomic investigations focus on the averaged metabolism of millions of cells and do not reflect fluctuations at the cellular or subcellular level, but organelle or “spatial” metabolomics can detect subcellular abnormalities induced by a disease or treatment. One challenge in understanding the impact of altered metabolism induced by IDH1^*mut*^ arises from our inability to determine compartment-specific metabolism (Wellen and Snyder, 2019).

Although significant progress has been made in metabolomics methods, to date, no technology is capable of addressing the spatial metabolomics problem in live cells (Rappez et al., 2019, Geier et al., 2019, Qi et al., 2018, Duncan et al., 2019, Ibanez et al., 2013, Lee et al., 2019). To address these challenges, we recently developed an automated Raman micro-spectroscopy approach, which allows us to quantify and monitor biomolecular composition in single organelles of live cells (Kuzmin et al., 2018, Lita et al., 2019). In the context of lipidomics, this approach facilitates a) measuring total lipid accumulation, b) identifying phospholipids and sterols, and c) partially characterizing the structure of lipids, including the degree of unsaturation and the ratio between cis and trans isoforms. We applied this novel method to investigate the organelle-specific metabolic alterations that occur as a result of IDH1^*mut*^ overexpression in a model of U251 glioblastoma cells. Our Raman microscopy–based spatial metabolic profiling revealed an increase in heterogeneity in lipid distribution across all organelles and an increase in lipid unsaturation in the ER due to IDH1^*mut*^ overexpression. Follow-up liquid chromatography (LC)/MS-based organelle lipidomics identified higher levels of saturated fatty acids (SFAs) and monounsaturated fatty acids (MUFAs) in the ER, which were subsequently incorporated into phospholipids (phosphoethanolamines and phosphocholines) of the ER membrane. To gain insight into the effects of these alterations on organelle function, we performed confocal and transmission electron microscopy (TEM). Our analyses revealed overall organelle dysfunction, as well as unique dilatation and membrane fragmentation in Golgi apparatus attributable to the IDH1^*mut*^; they were recapitulated in tumor tissue from patients and point to vulnerabilities that can be exploited therapeutically.

## Results

### IDH1 mutations induced heterogenous lipid composition in organelles

We recently developed a Raman spectroscopy method to selectively detect and quantify major types of biomolecules within single organelles in live cells (Kuzmin et al., 2018, Lita et al., 2019). Deconvolution of Raman spectra by *BCAbox* software allowed us to analyze lipid vibrational bonds that belong to different lipid species (Figure 1a, 1b). To understand the role of IDH1 mutation in lipid metabolism, we used this approach to profile cells with wildtype IDH1 (IDH1^*WT*^) and cells with the R132H or R132C IDH1 mutations (Liu et al., 2019), both of which generate different concentrations of D-2HG (Figure 5c, Supplementary Figure 1a). The Raman-based findings in live cells were complemented by conventional untargeted metabolomic analyses of isolated organelles. Raman spectral analysis showed that introducing the IDH1 mutation into a U251 glioblastoma cell line (U251^*R132H/C*^) increased lipid heterogeneity, as measured by the following parameters: 1) the lipid unsaturation parameter (the double bond contents, LSU) (Figure 1c), 2) the trans/cis parameter (stereoisomers of fatty acids) (Figure 1d) and 3) sphingomyelin and cholesterol levels (Figure 1e–f). In all organelles except lysosomes, the heterogeneity in distribution of lipid species decreased after addition of AGI-5198, a specific inhibitor of the IDH1 mutation (Figure 1g, Supplementary Figures S1 and S2). Protein content was more heterogeneous in Golgi apparatus of U251^*R132H*^ compared with U251^*WT*^ cells (Figure 1h). Averaged analyses of the biomolecular composition of single organelles revealed significant accumulation of total lipids in Golgi apparatus, mitochondria, and lysosomes in U251^*R132H/C*^ cells and the specific accumulation of sphingomyelin in ER, Golgi apparatus, and lysosomes of mutated cells (Figure 1i and 1j). Averaged RNA/DNA content was lower in the ER and mitochondria of U251^*R132H*^ cells compared with U251^*WT*^ cells. Total protein content was higher in ER in both U251^*R132C*^ and U251^*R132H*^ cells (Supplementary Figure 1b).

**Figure 1.**
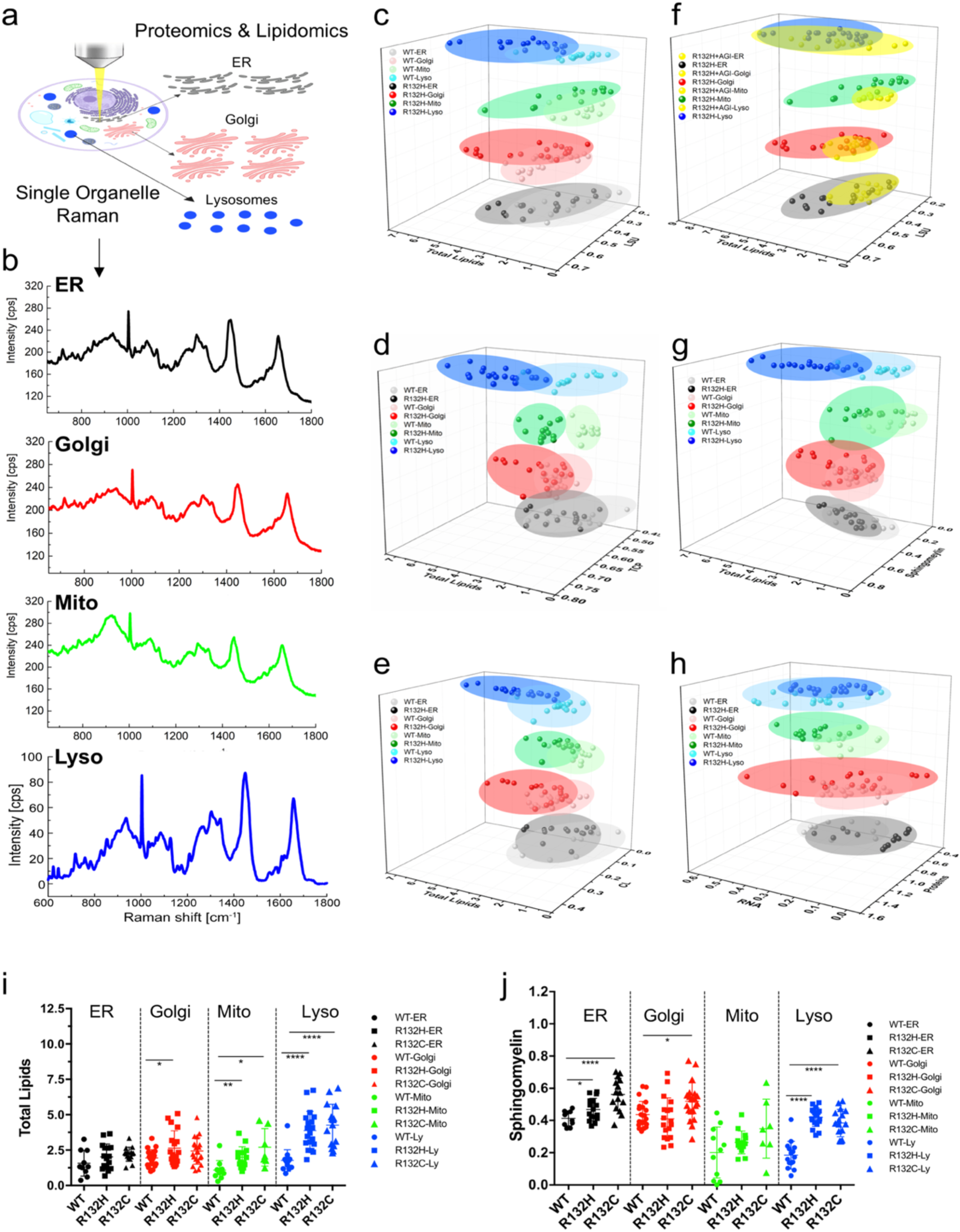
Global biomolecular changes induced by IDH mutation in live cells at the organellar level. a) Schematic representation of the strategy for the study. b) Representative Raman spectra of live cells obtained using our newly developed method (black, ER; red, Golgi; green, mitochondria; blue, lysosomes). c–f) Distribution of lipid unsaturation parameter in U251^*WT*^ and U251^*R132H*^ cells, represented for each cell and organelle to show the heterogeneity in lipid distribution across cells and organelles. Light ovals depict U251^*WT*^ organelle data; dark ovals, the U251^*R132H*^ data. g) The distribution of the lipid unsaturation parameter becomes more homogeneous after addition of AGI-5198, the inhibitor of IDH1 mutation (yellow ovals). h) Proteins and RNA values are heterogeneously distributed in U251^*R132H*^ cells as well. i and j) Averaged total lipid and sphingomyelin levels for each organelle depicting the changes as a function of the mutations.

### IDH1 mutations led to specific damage of the mitochondrial membrane

To gain more insight into the link between the unique lipid profile of U251^*R132H/C*^ cells and its cellular function, we compared the parameters obtained from the Raman measurements with the structure of each organelle obtained by Transmission Electron Microscopy (TEM). Using peak intensities at 1440 cm^-1^ and 1660 cm^-1^ from the organelle-specific Raman spectra, we extracted a parameter called the lipid unsaturation parameter (LSU), which quantifies the degree of unsaturation in lipids (Supplementary Figure 1c). The LSU parameter was significantly lower in the mitochondria of U251^*R132H*^ cells compared with those of U251^*WT*^ cells and was partially restored by the addition IDH1 mutation inhibitor, AGI-5198. Mitochondrial lipid unsaturation did not differ significantly between U251^*R132C*^ and U251^*WT*^ cells, according to the LSU parameter (Figure 2a). A lower LSU parameter value suggests greater content of lipids with saturated bonds (C-C) according to our calibration with fatty acid standards (Figure 2b). The trend in LSU parameter correlates with a trend observed for partial damage of mitochondria in these cells. Although 85% of the mitochondria of U251^*R132H*^ cells were partially damaged, only 50% of mitochondria in U251^*WT*^ and U251^*R132C*^ cells were damaged. Addition of the IDH1 mutant inhibitor AGI-5198 restored the number of partially damaged mitochondria in U251^*R132H*^ cells to the same level as in U251^*WT*^ cells (Figure 2c). Unexpectedly, mitochondria of U251^*WT*^ and U251^*R132H*^ cells were damaged differently. Whereas the U251^*WT*^ cells had both inner and outer mitochondrial membrane damage with more pronounced outer membrane damage, U251^*R132H*^ cells showed more inner mitochondrial damage that led to cristolysis and matrix lysis (Figure 2d-g). These differences were also noticeable in tissue from five patients with either IDH^*WT*^ glioblastoma multiforme or IDH^*mut*^ oligodendroglioma (Figure 2h-n). Tissue from patients with IDH^*mut*^ oligodendroglioma had round mitochondria and more pronounced inner membrane defects leading to cristolysis (Figure 2i-k), whereas tissue from IDH^*WT*^ glioblastoma multiforme (GBM) or the tumor margin displayed both round and elongated mitochondria with more pronounced outer membrane defects (Figure 2l-n).

**Figure 2.**
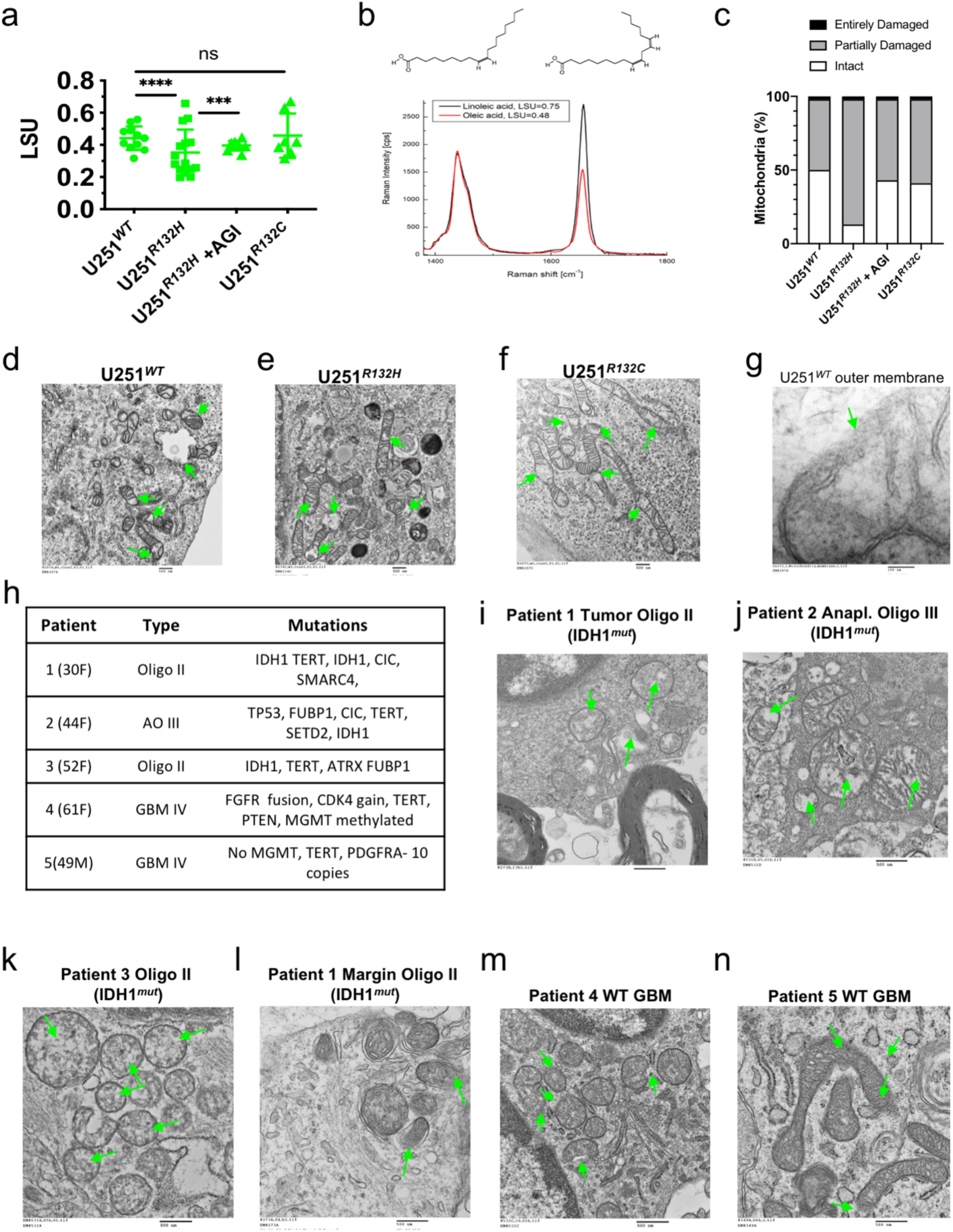
IDH1 mutation induces lower mitochondrial LSU parameter and inner membrane mitochondrial damage. a) The lipid unsaturation (LSU) parameter was significantly lower in U251^*R132H*^ cells and partially recovered after the inhibitor AGI-5198 was added. b) LSU parameter varies from close to 0.5 in the case of oleic acid (one C=C) to 0.75 for linoleic acid (two C=C bonds). c) Introduction of an R132H mutation to U251 cells resulted in more partially damaged mitochondria (80% of cells), whereas addition of the inhibitor AGI-5198 resulted in recovery of mitochondria to 50% of cells, similar to U251^*WT*^ and U251^*R132C*^. d–f) Representative electron micrographs of U251^*WT*^, U251^*R132H*^, and U251^*R132C*^. Green arrows indicate damaged regions of the mitochondria. g) Transmission electron micrograph of outer membrane breakage in U251^WT^ cells. h) The molecular characterization and histology of patient samples used in this study. i–k) Transmission electron micrographs show inner mitochondrial damage specific to oligodendroglioma tissue. Green arrows indicate the loss of inner membrane integrity. l–n) Transmission electron micrographs show outer mitochondrial damage in IDH1^*WT*^ glioblastoma tissue as indicated by the green arrows.

### IDH1 mutations led to an increased number of lysosomes

Using Raman analysis, we found significant upregulation of lipid and sphingomyelin content in lysosomes as a function of both mutations (Figure 1i, 1j). To understand the consequences of increased lipid composition we then performed both TEM and confocal microscopy of cells. TEM identified an increased number of lysosomes in both U251^*R132H*^ and U251^*R132C*^ cells compared with U251^*WT*^; this finding was further validated by LAMP-1-based western blot analysis (Supplementary Figure 3a, 3c, and 3h). Lysosomal area was greater only in U251^*R132H*^ cells, but not U251^*WT*^ cells (Supplementary Figure 3i). Lysosomes had higher electron density in mutated cells when analyzed via TEM, and this was recapitulated in lysosomes from patient samples of IDH^*mut*^ oligodendroglioma (Supplementary Figure 3a, 3d and 3e). Addition of the IDH1 mutant inhibitor AGI-5198 to all U251 cells led to a significant increase in total lysosomal lipid content, lysosome number, and their trafficking speed (Supplementary Figure 3j-m, Movies 1 and 2). TEM images of cells treated with AGI-5198 also showed increased lamellar content in these lysosomes, reflective of phospholipid accumulation (Supplementary Figure 3d). These results suggest that introduction of IDH1^*mut*^ leads to increased number of lysosomes, their function, and increased phospholipid accumulation in the lysosomes.

**Figure 3.**
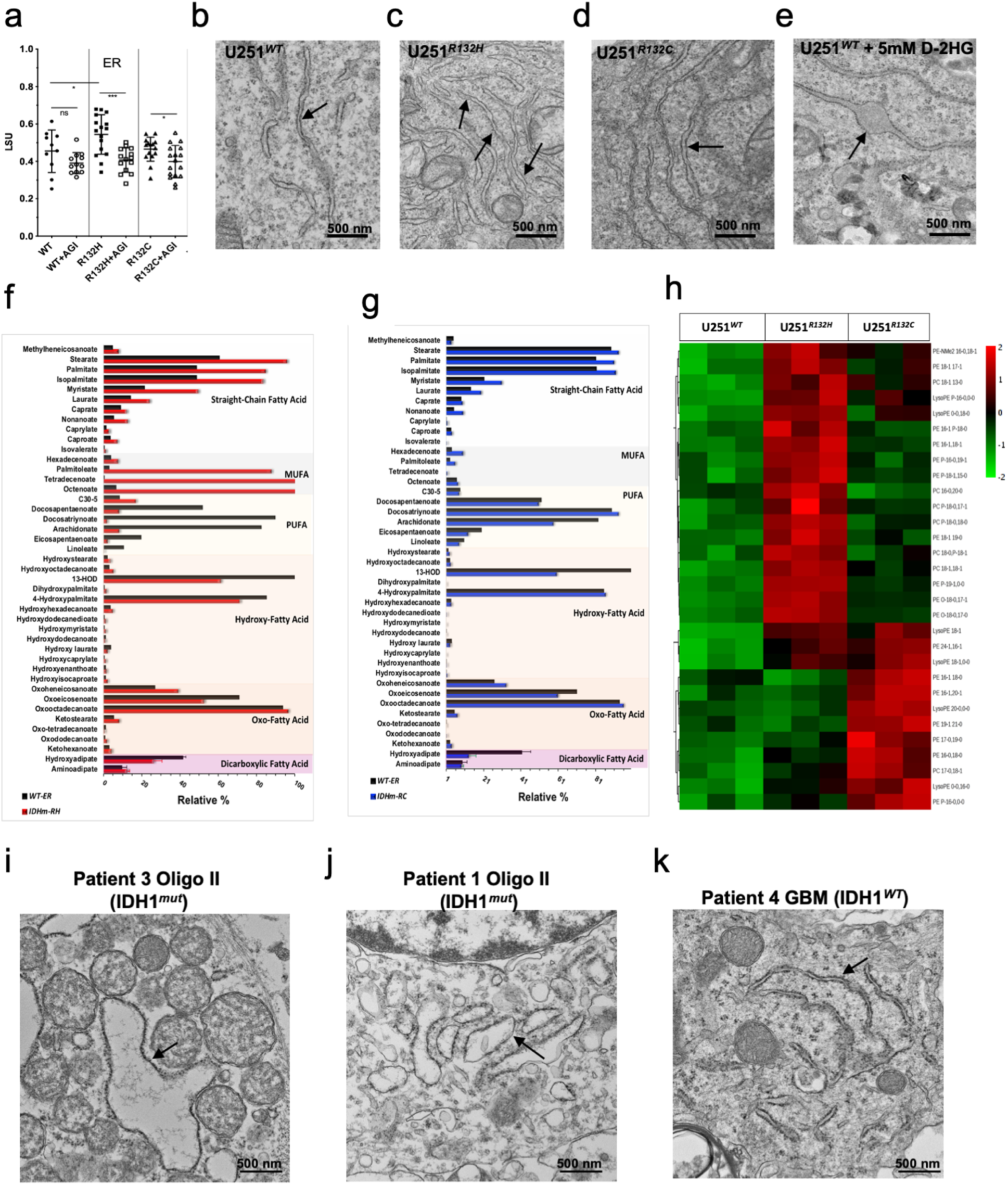
Endoplasmic reticulum (ER)-specific lipid changes due to IDH1 mutation. a) Lipid unsaturation (LSU) parameter from Raman analysis shows higher LSU in the ER of U251^R132H^ cells only. b–d) Transmission electron micrographs of U251^*WT*^, U251^*R132H*^ and U251^*R132C*^ show no changes in the ER structure. Arrows indicate ER. e) Addition of 2HG to U251^*WT*^ leads to ER dilation. f) and g) Comparison of the lipidomic profiles of ER from U251^*WT*^ (black) and U251^*R132H*^ (red) or U251^R132C^ (blue) revealed a higher percentage of saturated and monounsaturated fatty acids in U251^R132H^ cells only, similar to the LSU parameter. Black bars represent U251^*WT*^ cells while the red and blue represent U251^*R132H*^ and U251^*R132C*^, respectively. g) Heatmap of phospholipids extracted from the ER of U251^*WT*^, U251^*R132H*^, and U251^*R132C*^ cells shows the 30 most significantly altered features. h–j) Tissue from patients with IDH^*mut*^ oligodendroglioma showed dilated ER, but tissue from a GBM patient (IDH^*WT*^) did not.

### IDH1 mutation leads to upregulation of saturated or monounsaturated fatty acids and phospholipids containing these species in the endoplasmic reticulum

The LSU parameter was significantly higher in U251^*R132H*^, but not statistically significant U251^*R132C*^ compared with U251^*WT*^ in the ER and was restored upon addition of IDH1-mutant inhibitor, AGI-5198 (Figure 3a). A higher LSU parameter suggested more lipids with C=C bonds (Figure 3b, Supplemental Figure 1c). Interestingly, the TEM images of the ER do not show any differences between U251^*WT*^ and U251 ^*R132H/C*^ cells (Figure 3b, 3c and 3d) however, addition of 5 mM D-2HG leads to ER dilation (Figure 3e). Because the LSU does not discriminate between the types of lipids and the number of contributing C=C bonds, we conducted a lipidomic profile of isolated ER organelles from U251^*WT*^ and U251^*R132H/C*^ glioma cells using LC/MS (Figure 3e, 3f). ER-specific lipidomic analysis showed a higher percentage of SFAs and MUFAs and a lower percentage of polyunsaturated fatty acids (PUFAs) in U251^*R132H*^ compared to IDH^*WT*^ cells (Figure 3f). U251^*R132C*^ cells did not show the same striking differences, just minor upregulation of SFAs, which coincides with no change in the LSU parameter obtained from the Raman measurements (Figure 3g). The abundant saturated and monounsaturated saturated fatty acids incorporated into the phospholipids from ER in both mutant cells (Figure 3h). Interestingly, samples from oligodendroglioma (IDH^*mut*^) displayed vastly dilated rough ER (Figure 3i, 3j), which was not evident in GBM tissue samples (Figure 3k).

### IDH1 mutation leads to depletion of SFA-, and MUFA-PEs and PCs from Golgi apparatus

Untargeted organelle lipidomics via MS revealed that elevated SFAs and MUFAs in the ER was correlated with higher phospholipid levels in ER that contained those lipid species in U251^*R132H/C*^ cells specifically (Figure 3g). Our analysis also revealed downregulation of SFA and MUFA phospholipids in Golgi apparatus of these cells as a result of both IDH1 mutations (Figure 4a). Further, TEM studies showed that Golgi cisternae were enlarged and swollen in IDH1^*R132H*^ cells. The swollen and dilated stacks of Golgi observed in U251^*R132H*^ cells were restored to normal after adding AGI-5198 (Figure 4b). In this study, the drastic dilation of Golgi cisternae that was more pronounced in U251^*R132H*^ than U251^*R132C*^ cells was further investigated. First, to confirm the deregulation of PEs and PCs at the organelle level due to IDH1 mutation in live cells, we stained the ER Golgi, mitochondria and lysosomes with red fluorescence protein (RFP)-proteins and added BODIPY™ FL C16 (4,4-Difluoro-5,7-Dimethyl-4-Bora-3a,4a-Diaza-s-Indacene-3-Hexadecanoic Acid) to visualize the uptake and tracing of this SFA’s fate by using confocal microscopy. Although BODIPY-palmitate appears to be ubiquitously distributed throughout the cytoplasm, we observed no uptake of BODIPY-palmitate in the ER, mitochondria or the lysosomes by the lack of colocalization with the resident membrane red fluorescent protein in U251^*R132H/C*^ or U251^*WT*^ cells (Supplementary Figure 4a-c). No change in this distribution was observed after addition of the IDH1^*mut*^ inhibitor, AGI-5198 (Supplementary Figure 4a-c bottom panels). Prominent colocalization of BODIPY-palmitate with the Golgi-resident protein N-acetylgalactosaminyltransferase was observed specifically in U251^*R132H*^ cells; this effect could be reversed by adding AGI-5198 (Figure 4c, 4d). Specific uptake of SFA by Golgi organelles indirectly suggests the lack of SFA in Golgi and correlates with the depleted PEs observed via MS-based lipidomics. Indeed, the most downregulated features, as seen in the heat map (Figure 4a) or the volcano and bar plots (Figure 4e, 4f) when U251^*R132H/C*^ was compared with U251^*WT*^, were the SFA- and MUFA-based PEs and PCs, which were partially restored by adding AGI-5198 (Figure 4f).

**Figure 4.**
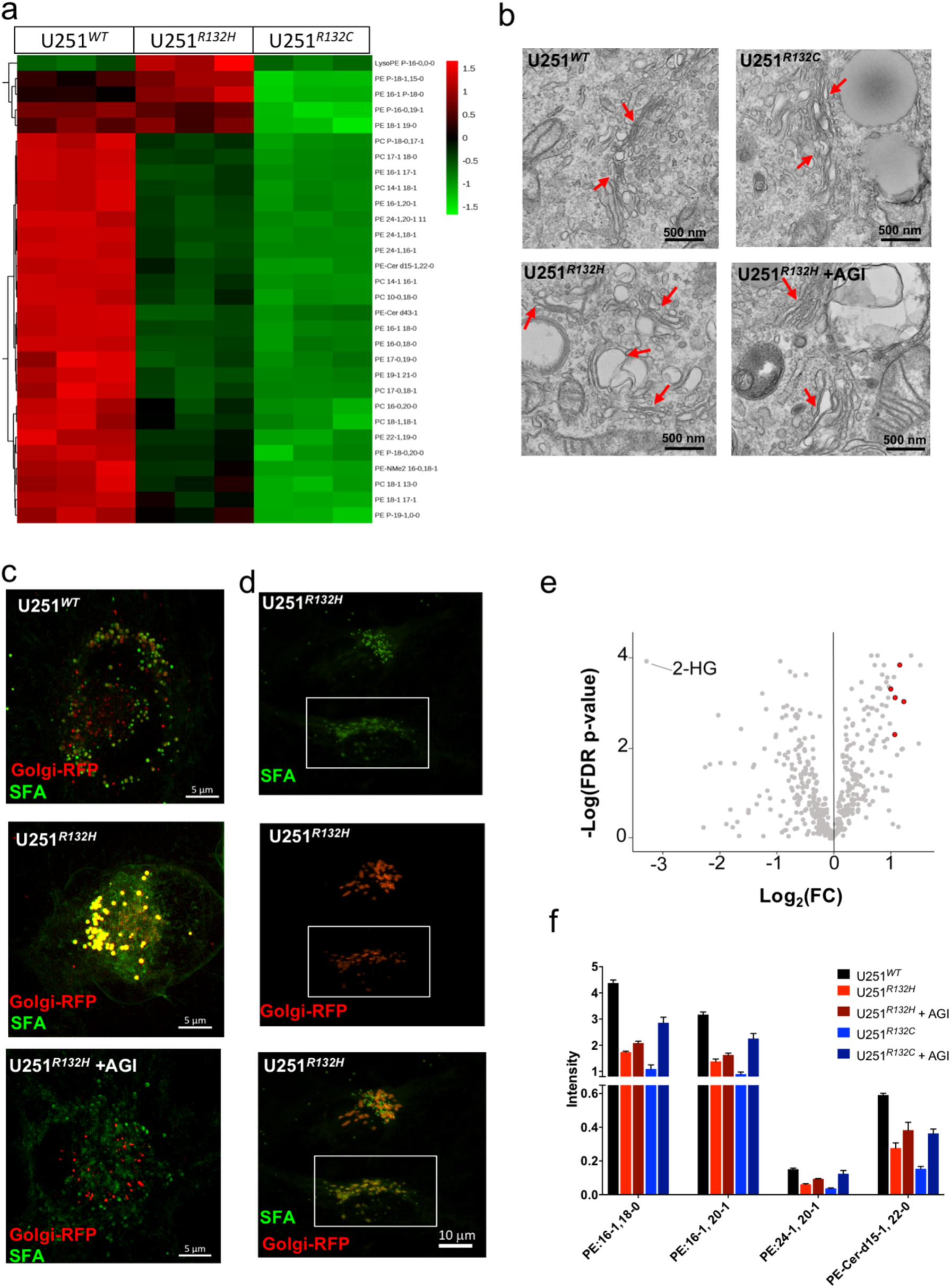
Depletion of phospholipids in Golgi apparatus is correlated with Golgi dilation. a) Heatmap with the most significant phospholipids from the untargeted Golgi lipidomics showed that most phospholipids containing saturated or monounsaturated phospholipids are depleted in Golgi. b) Transmission electron micrograph shows altered Golgi structure in U251^*R132H*^ and to a lesser extent in U251^*R132C*^ cells. Addition of AGI-5198 partially restored the Golgi structure in U251^*R132H*^ cells. c) Fluorescence microscopy shows co-localization (yellow, middle panel) of saturated fatty acids (green) with the Golgi apparatus (red) in U251^*R132H*^ cells and the loss of co-localization in the presence of the inhibitor AGI-5198. d) Z-stacked Golgi reconstructed image shows the area of co-localization. e) Volcano plot of the lipidomic assay comparing U251^*WT*^ and U251^*R132H/C*^ combined. 2-HG appears to be the most significant metabolite upregulated in the mutant cells, whereas phospholipids (PEs) (red dots) are among the most downregulated lipids in mutant cells. f) Relative intensity of PEs that contain zero or one double bond are downregulated in mutant cells (light red and light blue bars) compared with wildtype (black bars) and are partially restored by adding AGI-5198 inhibitor (dark red and dark blue bars).

**Figure 5.**
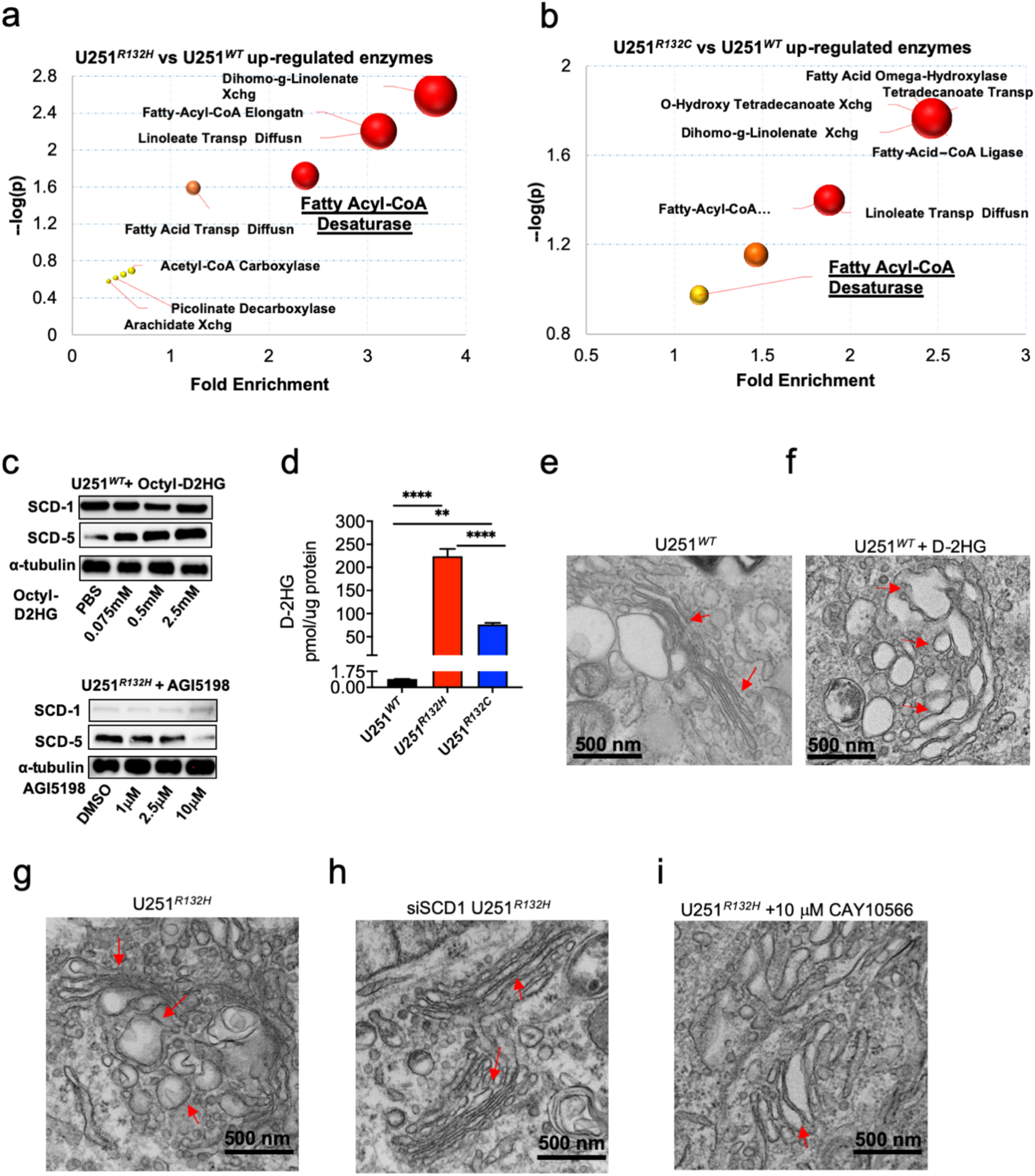
Stearyl Co-A desaturase (SCD) overexpression is induced by D-2HG and is responsible for IDH1^*mut*^-induced membrane defects in Golgi.. a) and b) Predicted enzymes from Golgi-specific lipids identified from U251^*R132H/C*^ by mass spectrometry. c) Western blot analysis show upregulation of SCD-5 in mutant cells and upon addition of 0.5-2.5 mM D-2HG and the decreased with the addition of AGI5198. d) Compared with U251^*WT*^, D-2HG concentration is 240-fold higher in U251^*R132H*^ and 90-fold higher in U251^*R132C*^ cells. e) and f) Addition of D-2HG in U251^*WT*^ was enough to cause Golgi dilation. g) and h) Comparison between Golgi of U251^*R132H*^ cells and the same cells that lack SCD-1 showed a restored Golgi structure in the cells lacking SCD-1. i) Inhibiting the SCD-1 enzyme with CAY10566 led to partial restoration of Golgi structure.

### SCD enzyme is responsible for Golgi dilation

To understand the link between mutant IDH1*-*induced imbalance in lipid distribution and the organellar defects described above, we performed enzyme prediction from the metabolomics on the affected organelles. We used U251^*R132H/C*^ Golgi metabolites to perform untargeted lipidomic analysis of Golgi apparatus followed by enzyme prediction. We found that desaturases, hydrolases, and lipid transport were the major upregulated enzymes in the mutated cells (Figure 5a, 5b). The desaturases in U251^*R132H*^ cells correlated more significantly with metabolite trends than did those in U251^*R132C*^ cells. Western blot analysis showed greater expression of SCD1 in U251^*R132H*^ and U251^R132C^ cells and minimal expression of SCD1 in U251^*WT*^ (Figure 5c). SCD enzymes utilize Fe^2+^, NADPH, cytochrome b5, and O_2_ to catalyze the first step in MUFA biosynthesis from saturated fatty acids. Using The Cancer Genome Atlas data (TCGA), we found that SCD enzymes were also upregulated in an unsupervised cluster analysis done on all the mRNA levels from the fatty acid synthesis pathway (Supplementary Figure 5c), suggesting that those genes are important in these patient samples.

### Addition of D-2HG increased SCD-5 expression and Golgi dilation, whereas knockdown of SCD1 restored the Golgi structure in U251^*R132H*^

Because U251^*R132H*^ and U251^*R132C*^ cells differ in the level of D-2HG that they produce (Figure 5d), we next tested whether addition of D-2HG to WT U251 cells is sufficient to induce the dilated Golgi phenotype observed in U251^*R132H*^ cells. Addition of 5 mM D-2HG to U251^*WT*^ cells induced that phenotype (Figure 5e, 6f). We then confirmed the hypothesis that D-2HG is sufficient to upregulate the SCD enzymes. Indeed, western blot analysis showed that D-2HG incubation increased expression of SCD5 enzyme, while not affecting SCD1 expression (Figure 5c). However, knockdown of SCD1 via short hairpin RNA restored the Golgi structure in U251^*R132H*^ cells, as measured by TEM (Figure 5g and 5h). Addition of an SCD inhibitor also restored the Golgi of U251^*R132H*^ cells (Figure 51). Inhibition of D-2HG via the AGI-5198 inhibitor also restores Golgi morphology in U251^*R132H*^ cells (Figure 4b). Together these results support a working model in which D-2HG induces SCD overexpression, which leads to Golgi defects and apoptosis.

### SCD overexpression was associated with Golgi dilation and longer survival in oligodendroglioma patients

Using gene expression data and survival data available through The Cancer Genome Atlas (TCGA), we compared the expression of the both SCD isoforms (SCD-1 and SCD-5), the first enzyme in MUFA biosynthesis. Consensus clustering of mRNA for fatty acid synthesis genes, revealed strikingly high expression of both SCD-1 and SCD-5 in oligodendroglioma tissues (Figure 6a). The IDH^*mut*^ low-grade gliomas had significantly higher mRNA levels for SCD-1 and SCD-5 enzymes than IDH^*WT*^ tumors (Figure 6b). We also compared mRNA expression across all histological types and grades of glioma and found that similar to the heat map, patients with oligodendroglioma had the highest mRNA levels of SCD-1 and SCD-5, followed by those with astrocytoma and GBMs (Figure 6c). Kaplan-Meyer overall survival analysis from TCGA database revealed a significant association between SCD-1 and SCD-5 expression levels and survival in patients with IDH1^*mut*^ oligodendroglioma. (Figure 6d, 6f). Patients with higher levels of SCD-1 and SCD-5 (SCD^*High*^) had longer survival than patients with lower levels of SCD1 and SCD-5 (SCD^*Low*^); this benefit was not seen in patients with other subtypes of gliomas (Supplemental Figure 5a-h) or IDH^*WT*^.

**Figure 6.**
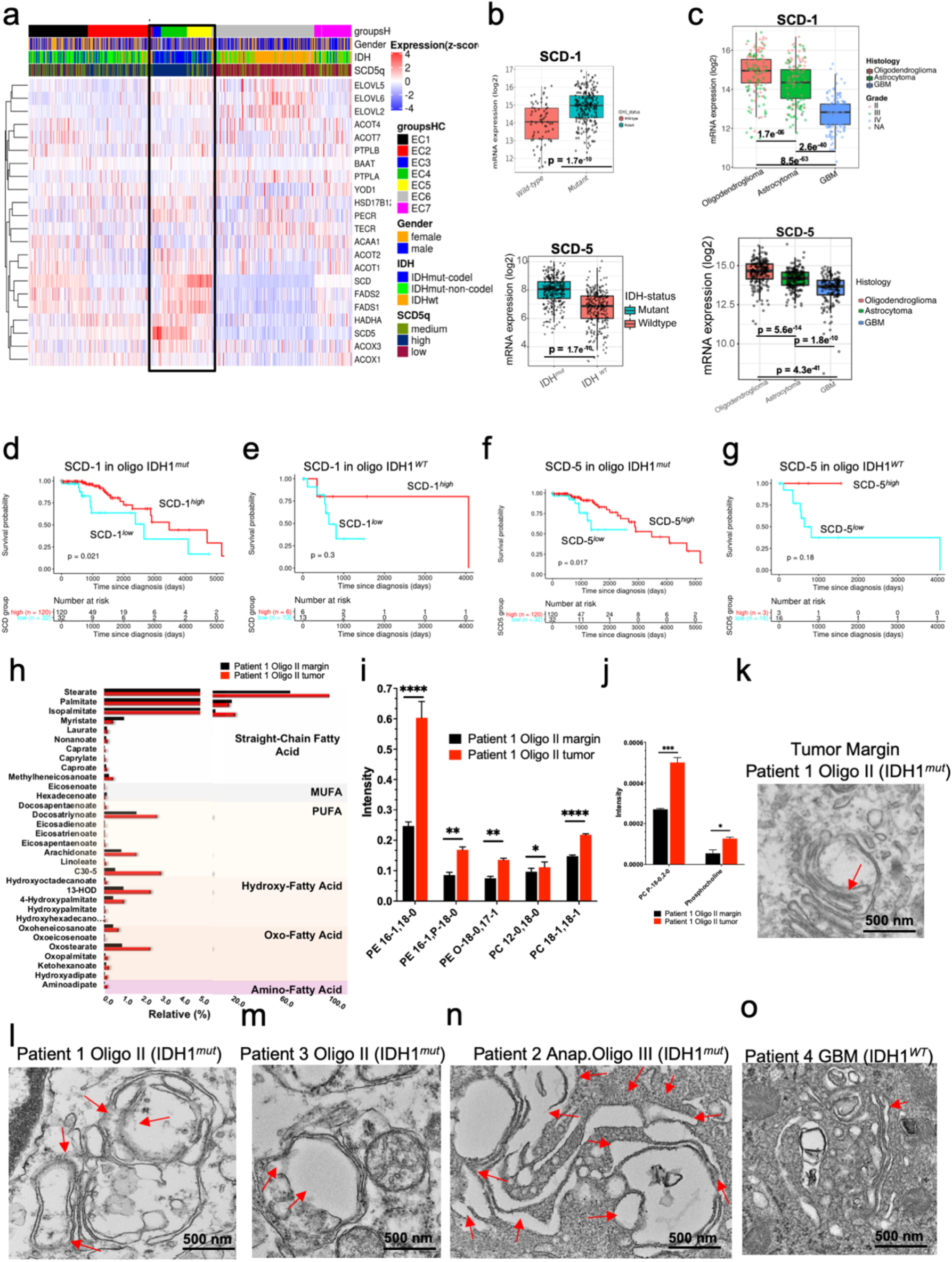
Golgi-specific phospholipid imbalance and Golgi dilation are prevalent in an oligodendroglioma patient sample and are correlated with stearyl Co-A desaturase (SCD-1 and SCD-5) overexpression and longer survival. a) Consensus clustering of 22 genes from fatty acid synthesis pathway reveled clustering of oligodendroglioma samples with highest mRNA levels for SCD-1 and SCD-5 enzymes. b) mRNA levels of SCD-1 and SCD-5 from The Cancer Genome Atlas show increased expression in these transcripts in IDH1^*mut*^ tissue compared with IDH ^*WT*^ in low-grade gliomas (LGG). c) mRNA levels of SCD-1 and SCD-5 were inversely correlated with the molecular subtypes in the following order: oligodendroglioma, astrocytoma, and GBM. Figure was created using GlioVis data portal (Bowman et al., 2017). d–g) SCD-1^*high*^ and SCD-5^*high*^ expression were correlated with better survival only in IDH1^*mut*^ oligodendroglioma, however, and no survival benefit was observed in IDH1^WT^ oligodendroglioma. h-j) Increased SFA- and MUFA- and its phospholipids in the Golgi of a tumor from patient 1 correlates with the in vitro data. k–o) Transmission electron micrographs of Golgi in tissue from different grades of oligodendroglioma compared with glioblastoma multiforme (GBM) (IDH1^WT^) show specific dilation of this organelle (red arrows) in the oligo tumor samples only.

To understand the clinical relevance of our *in vitro* findings, and recapitulate the TCGA findings, we obtained fresh tumor from patients with either IDH1^*WT*^ GBM or oligodendroglioma (IDH1^*mut*^) with the molecular characteristics shown in Figure 2h. We confirmed that tissue from oligodendrogliomas contained both higher MUFAs and PE-MUFAs. We extracted the Golgi apparatus from tissue of Patient 1 as well as tumor margin and performed untargeted lipidomics via LC/MS. Golgi apparatus from this tumor contained higher levels of MUFAs as well as PC- and PE-MUFAs compared with the margin from the same patient (Figure 6h-j). We then confirmed that Golgi apparatus was normal for the tumor margin, while dilated and swollen in the tumor sample (Figure 6l). Additionally, we measured other oligodendroglioma tissues and found dilated Golgi, while GBM tissue showed close to normal Golgi (Figure 6m-o). Thus, our TEM analysis of three IDH1^*mut*^ (oligodendroglioma) patient samples and two IDH^*WT*^ (glioblastoma) tissue samples revealed that IDH1^*mut*^ tumors had the same structural defects in the Golgi apparatus as those found *in vitro*, whereas the IDH^*WT*^ samples did not have the same level of dilation compared with the IDH1^*mut*^.

### Increasing MUFAs levels leads to further Golgi’s cisternae dilation, apoptosis, and cell death in IDH^*mut*^ cell lines

Next, we studied the consequences of tilting the balance toward more MUFA levels by adding oleic acid, the product of the SCD1 enzyme. TEM micrographs showed Golgi cisternae dilation after adding oleic acid (Figure 7a-c). Oleic acid changed the morphology of the cells: vesicles resembling lipid droplets appeared inside the cells (Figure 7d, 7e). Oil red and neutral lipid staining (LipidTOX Neutral Green, Thermofisher) confirmed the formation of lipid droplets (Supplementary Figure 6g). Using a panel of IDH1^*R132H*^ patient-derived cell lines from astrocytoma (BT142, NCH1681), as well as oligodendroglioma (TS603) and GBM (GSC923, GSC827), we confirmed that SCD1 is highly expressed in patient-derived IDH1^*mut*^ cells, with the highest amount of SCD1 in oligodendrogliomas, whereas basal expression of SCD1 was observed in IDH1^*WT*^ patient-derived neurospheres (Supplementary Figure 6a, 6b). With this panel, we next explored whether we could manipulate the specific vulnerability induced by IDH1 mutation for therapeutic purposes. Since the most prevalent MUFAs in the cells are palmitoleic and oleic acid, we added oleic acid to U251^*R132H/C*^ and patient-derived IDH1^*mut*^ and IDH1^*WT*^ cell lines, as described above. Cell proliferation and viability decreased significantly within the first 24 hours after 100 μM oleic acid was added, as revealed by trypan blue viability assays (Figure 7f-h, Supplementary Figure 6c). The same concentration in IDH^*WT*^ cell lines did not alter their proliferation rate or viability as drastically as it did for IDH1^*mut*^ cells (Figure 7i). IDH1^*mut*^ cells were more sensitive to oleic acid–induced apoptosis, as measured by EC_50_ using a cell viability kit CCK-8 assay and Annexin V and a 7-AAPC flow cytometry–based assay. All patient-derived IDH1^*mut*^ cell lines irrespective of their molecular type (oligodendroglioma or astrocytoma) were 4-fold to 7-fold more sensitive to oleic acid–induced cell death, respectively, as measured by EC_50_, compared with WT (GSC 827) cells (Figure 7j, 7k). Addition of oleic acid increased the percentage of cells that underwent late apoptosis, via flow cytometry (43% apoptotic cells in IDH1^*mut*^ compared with 7% in IDH^*WT*^). Inhibition of fatty acid synthase (FASN) or addition of PUFAs (linoleic acid) had a more pronounced effect on U251^*WT*^ cells but did not affect U251^*R132H/C*^ cells significantly, suggesting that the specific vulnerability of IDH^*mut*^ is in the accumulation of MUFAs and SFA-to-MUFA conversion (Supplementary Figure 6e, 6f).

**Figure 7.**
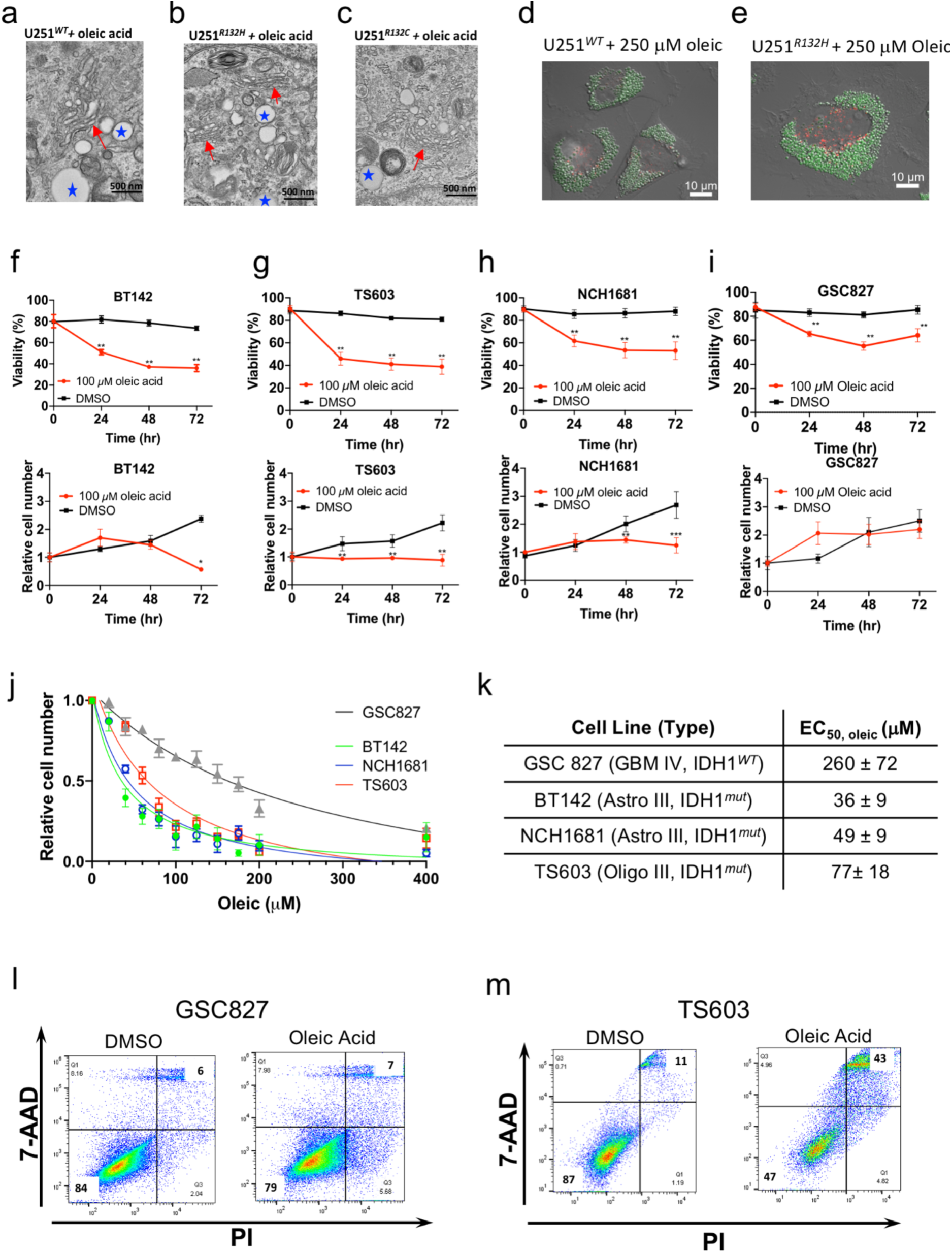
Addition of MUFA leads to IDH1^*mut*^-specific cell death. a–c) Addition of oleic acid in U251^*WT*^ and U251^*R132H/C*^ cells increased Golgi dilation. d) and e) Addition of oleic acid led to the accumulation of lipid droplets followed by cell death via apoptosis. f–i) Oleic acid treatment also led to decreased viability in patient-derived cell lines BT142, TS603, and NCH1681, and it was more pronounced than in IDH^*WT*^ (GSC827). j) and k) Cell counting, CCK-8 assay shows greater sensitivity of IDH1^*mut*^ cells (green, blue, and red lines) to oleic acid supplementation compared with IDH^*WT*^ neutrospheres (gray line). l) and m) In IDH^*WT*^ cells (GSC827), there was no change in cellular apoptosis measured 48h hours after addition of oleic acid. In patient derived IDH^*mut*^ cells (TS603) 43% of cells underwent late apoptosis.

### Organelle proteomics revealed the link between D-2HG and SCD expression

To understand the mechanism by which D-2HG induced SCD specific overexpression, we conducted organelle specific proteomics. Interestingly, FTL and FTH were upregulated in the Golgi proteome of IDH^*mut*^ oligodendroglioma tissue and U251^*R132H*^ cells compared with either margin or U251^*WT*^, respectively (Figure 8a, 8b, 8c). Reactome and Gene Ontology databases suggested a direct link between FTL and FTH1 and Golgi vesicle, biogenesis, budding and transport (Figure 8d). Western Blot analysis revealed the highest expression of FTL in TS603, the oligodendroglioma cell line as well as U251^*mut*^ cells (Figure 8e). Interestingly, deferoxamine addition led to inhibition of cell growth specifically for IDH1^*mut*^ cell lines compared with IDH^*WT*^ (Figure 8f, Supplementary Figure 7a, 7b).

**Figure 8:**
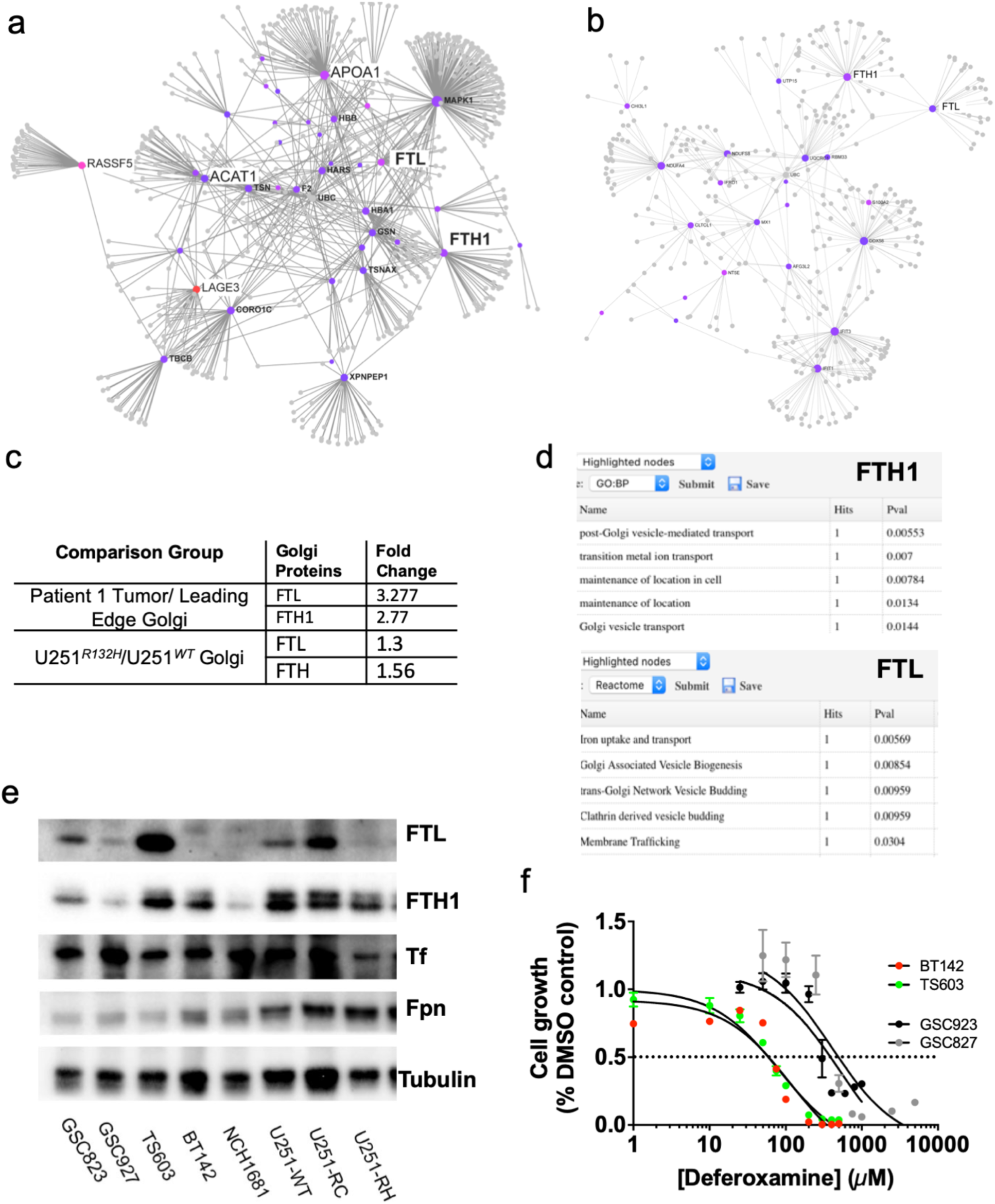
Ferritin is upregulated in Golgi proteome of tissue and cells. a) and b) Mass spectrometry-based proteomics analysis of Golgi proteins extracted from oligodendroglioma patient 1 and its margin as well as from U251^*R132H*^ or IDH^*WT*^ cells. c) Fold change of ferritin heavy chain (FTH1) and ferritin light chain (FTL) levels from Golgi tumors versus same organelle extracted from the margin or U251^*R132H*^ versus IDH^*WT*^ Golgi cells d) Reactome and Gene Ontology reveled significant pathways correlated with both FTH1 and FTL. e) Western Blot analysos of iron-based proteins in U251^*R132H/C*^ and IDH^*WT*^ as well as patient derived cell lines: BT142, TS603, NCH1681 (IDH1^*mut*^) and GSC 923 and GSC827 (IDH^*WT*^). f) Cell growth analysis of IDH^*WT*^ and IDH^*mut*^-patient derived cell lines as a function of increasing deferoxamine concentrations, an iron chelator, show increased sensitivity of IDH^*mut*^ cell lines.

## Discussion

Spatial and temporal compartmentalization of cellular metabolism is notoriously challenging to assess in part because of limitations in traditional MS-based whole-cell analysis (Wellen and Snyder, 2019). However, we could better understand localized metabolic alterations in diseases such as cancer by determining the distribution of metabolites in space and time. Recently, we developed a new method based upon Raman spectroscopy, coupled with fluorescence microscopy that allowed us to quantify classes of biomolecules at the organelle level (Lita et al., 2019, Kuzmin et al., 2018). Using this approach, here we quantified levels of proteins, lipids, DNA, RNA, cholesterol, and sphingomyelin at previously unmeasurable subcellular levels in a model system of IDH1^*mut*^ glioma.

Follow-up extraction of these organelles and MS-based lipidomics identified much higher percentages of saturated- and monounsaturated-phospholipids in the ER, which was complemented by lower percentages of those species of phospholipids in Golgi, specifically in U251^*R132H*^ cells. We confirmed the specific depletion of saturated and monounsaturated phospholipids by uptake experiments with BODIPY-palmitate, which preferentially co-localized in Golgi apparatus.

Because these phospholipids play an important role in membrane integrity, we next explored the link between the structure of organelles and phospholipid imbalance in the ER and GA. We used TEM to visualize the consequences of such alterations to membrane morphology in each organelle. We found global disruption of organelle integrity as a function of IDH mutation, which could be partially restored by addition of AGI-5198, an inhibitor of the IDH-mutant enzyme. In both cell lines and oligodendroglioma patient tissue, mitochondria lost internal cristae, a phenomenon known as cristolysis. These cristae are surrounded by an inner phospholipid bilayer that is rich in PEs; therefore, we speculate that a disturbed phospholipid (de Kroon et al., 1997) profile could affect their integrity. Because phospholipid composition of the outer membrane is very different from that of the inner mitochondrial membrane, it is tempting to speculate that MUFA-PEs are involved in such defects. Further investigations are needed to establish the role of altered phospholipids in mitochondrial structure and function. The number of lysosomes was higher in U251^*R132H/C*^ cells, which could indicate additional accumulated material resulting from organelle defects induced by IDH1^*mut*^ (Saftig, 2006). A nonspecific cytotoxic effect of AGI-5198 was also identified, as manifested by better lysosomal function in all cells in the presence of this inhibitor.

Golgi displayed swollen cisternae, which was confirmed in samples from patients with IDH1^*mut*^ and oligodendroglioma. Herein, we mainly explored the phospholipid distribution in the ER and Golgi. To narrow the search for enzyme(s) that might be responsible for the imbalance between SFAs, MUFAs, and PUFAs in the ER and Golgi, we conducted enzyme enrichment analysis using MetaboAnalyst and consensus clustering using public patient data from TCGA. Stearyl-CoA desaturase (SCD) was commonly altered in our analyses, with overexpression correlating with better survival of patients with oligodendroglioma (1p/19q co-deleted, IDH1^*mut*^). SCD is an integral membrane protein of the ER and an important enzyme in the biosynthesis of MUFA. It produces two common products—oleyl- and palmitoyl-CoA. Because oleic and palmitoleic acids are the major MUFAs in fat depots and membrane phospholipids, we explored the therapeutic consequences of tilting the balance toward more MUFAs in IDH1^*R132H*^ cells that were derived from patients or engineered to overexpress this mutation. We found that IDH1^*mut*^ cells were specifically vulnerable to MUFAs, whereas SFAs and PUFAs affected IDH1^*WT*^ cells. Addition of oleic acid decreased the viability and proliferation rate of all IDH1^*mut*^ cell lines derived from patient samples and, to a lesser degree, IDH1^*WT*^ cells, by causing cellular apoptosis. Oleic acid also caused massive intracellular lipid droplet accumulation and the formation of a foam-like cell morphology.

This imbalance in lipid composition was also evident in tissue from patients. The patient tissue analyzed in this study had the following characteristics: patients 1 and 2 only had surgery at the time of tissue analysis, whereas all other patients had the standard of care, which included radiation and chemotherapy. Tissue from patients 3 and 5 was from a recurrent tumor, and the others were from primary tumors. For patient 1, we profiled the Golgi specific lipids extracted from the tumor and margin and we identified greater PE-MUFA and PE-SFA in the tumor compared with the margin. Golgi cisternae were dilated in tissue from oligodendroglioma, but they were normal in tissue from GBM. These findings provide clinical relevance to our cellular studies by confirming that the defects exist in patient tissue.

To understand the mechanisms by which D-2HG increases the expression of SCD-5, we conducted proteomic analysis of extracted Golgi from either cell lines or tissue of oligodendroglioma. Proteomics reveled increased levels of FTL and FTH1 in tumors’ Golgi compared to the margin and this was recapitulated in our model systems of IDH^*mut*^ cells. The levels of FTL and FTH were confirmed by Western Blot analyses of the cells and were strengthen by the findings that IDH^*mut*^ cells derived from patients with oligodendroglioma displayed the highest sensitivity to deferoxamine, a known Fe chelator. Our working hypothesis is that D-2HG inhibition of α-keto-dependent dioxygenases releases high levels of Fe intracellularly, which accumulates. Fe accumulation leads to increased expression of SCD which causes the imbalance in MUFAs measured here.

Imbalances in SFAs, MUFA and PUFAs have been linked to several diseases, including cancer. One prominent feature of this imbalance is altered membrane fluidity; membranes with higher PUFA display greater fluidity. We hypothesize that the accumulation of SFAs and MUFAs leads to loss of membrane fluidity, resulting in less trafficking of proteins and lipids though those membranes and eventually membrane rupture. The imbalance in SFAs, MUFAs, and PUFAs appears specific to patients with IDH1^*mut*^ and has survival implications. Previous studies using NMR spectroscopy reported that total PE levels were reduced due to the IDH1 mutation; however, the composition of the fatty acids linked to the PEs was not studied(Viswanath et al., 2018). Herein, we identified differences in the chemical composition of the phospholipids, first with Raman microscopy and then by MS. This investigation demonstrates a functional link between upregulation of SCD enzyme, formation of SFA- and MUFA-PEs, and damaged Golgi in IDH1^*mut*^ gliomas. Such metabolic vulnerabilities could provide a more effective way to target essential pathways to which cancer cells are intrinsically susceptible.

## Acknowledgments

This work was supported by the National Institutes of Health’s Intramural Research Program, Center for Cancer Research, National Cancer Institute, and the FLEX Technology Development Award. This project was funded in whole with federal funds from the National Cancer Institute, National Institutes of Health, under contract HHSN26120080001E. A.K. is supported by the National Institute of General Medical Sciences of the National Institutes of Health under Award Number R44GM116193. The content of this publication does not necessarily reflect the views or policies of the Department of Health and Human Services, nor does mention of trade names, commercial products, or organizations imply endorsement by the U.S. Government. We thank Erina He at NIH Medical Arts, who did the graphical abstract.

## Author Contributions

AL and AP conducted the research ML supervised the research. AL, AP, AK, TD, TY, LZ, VRR, CB, NdV, ERN, RS, OC, MK, CHM, MRG, PP and ML contributed to the data acquisition, interpretation and writing of the paper.

