## Supplemental Methods and Figures for "IDH1 Mutations Induce Organelle Defects Via Dysregulated Phospholipids"

**Contact for Reagents and Resource Sharing.** For any material used in the study, please contact Mioara Larion at:

**KEY RESOURCES TABLE**

| REAGENT or RESOURCE | SOURCE | IDENTIFIER |
|---|---|---|
| **Antibodies** | | |
| SCD-1 | Abcam | ab19862 |
| FASN | Abcam | ab218306 |
| LAMP-1 | Abcam | ab25630 H4A3 |
| tubulin | Abcam | ab15568 |
| β-actin (D6A8) Rabbit | Cell signaling Technology | 8457 |
| SREBP1 | Abcam | ab28481 |
| **Chemicals, Peptides, and Recombinant Proteins** | | |
| Fatostatin A | Tocris | 4444 |
| Bodipy-C16 | ThermoFisher | D3821 |
| Halt™ Protease and Phosphatase Inhibitor Cocktail (100X) | ThermoFisher | 78442 |
| Palmitoleic Acid | Cayman Chemicals | 10009871 |
| Oleic Acid | Sigma-Aldrich | 75090-5ML |
| CAY10566 | Cayman Chemicals | 944808-88-2 |
| Linoleic acid | Sigma-Aldrich | L1012-1G |
| Cerulenin | Sigma-Aldrich | C2389-5MG |
| CellLight™ Golgi-RFP, BacMam 2.0 | ThermoFisher | C10504 |
| CellLight™ Lysosomes-RFP, BacMam 2.0 | ThermoFisher | C10504 |
| CellLight™ Mitochondria-RFP, BacMam 2.0 | ThermoFisher | C10601 |
| CellLight™ ER-RFP, BacMam 2.0 | ThermoFisher | C10591 |
| ER-Tracker™ Green(BODIPY™ FL Glibenclamide) | ThermoFisher | E34251 |
| Oil Red O solution | Sigma-Aldrich | O1391-250ML |

| | | |
|---|---|---|
| HCS LipidTOX™ Green Neutral Lipid Stain | ThermoFisher | H34475 |
| LysoTracker Red DND-99 | ThermoFisher | L7528 |
| Trilencer-27 Fluorescent-labeled transfection control siRNA duplex - 1 nmol | ORIGENE | SR30002 |
| SCD1 (SCD) Human siRNA Oligo Duplex | ORIGENE | SR304248 |
| SCD1 Human siRNA | Sigma-Aldrich | EHU108071-20ug |
| **Critical Commercial Assays** | | |
| Pierce™ BCA Protein Assay Kit | ThermoFisher | 23225 |
| MINUTE™ GOLGI APPARATUS ENRICHMENT KIT | Invent Biotechnologies | GO-037 |
| Minute™ ER Enrichment Kit | Invent Biotechnologies | ED-028 |
| Minute™ Lysosome Isolation Kit | Invent Biotechnologies | LY-034 |
| Cell Counting Kit-8 | Dojindo Molecular Tech. | CK04-13 |
| Annexin V PE Apoptosis Detection Kit | BD Biosciences | 559763 |
| FITC Annexin V Apoptosis Detection Kit with 7-AAD | BD Biosciences | 640922 |
| **Experimental Models: Cell Lines** | | |
| TS603 | MSKCC | N/A |
| BT142 | ATCC | ATCC® ACS-1018™ |
| GSC827 | NOB | N/A |
| GSC923 | NOB | N/A |
| U251 | NOB | N/A |
| NCH1681 | University of Heidelberg | N/A |
| **Software and Algorithms** | | |
| Prism | Graphpad | https://www.graphpad.com/scientific-software/prism/ |
| R | R Project for Statistical Computing | https://www.r-project.org/ |
| Metaboanalyst | McGill University | http://www.metaboanalyst.ca |
| MassHunter Quant | Agilent | |

| | | |
|---|---|---|
| Agilent Masshunter Profinder | Agilent | |
| Partek Genomic Suite | Partek | http://www.partek.com/introducing-partek-genomics-suite-version-70 |
| Thermo Scientific™ OMNIC™xi Software | ThermoFisher | https://www.thermofisher.com/order/catalog/product/IQLAADGABFFAHCMBDI |
| BCAbox Software | ACIS, LLC | http://acis-us.com/ |

**Materials and methods:**

***Cell models and culture***

TS603 (grade III oligodendroglioma), BT142 (grade III oligoastrocytoma) and NCH1681 (grade III astrocytoma) (Dettling et al., 2018) were grown in DMEM/F12 medium supplemented with 1% N2 growth supplement, heparin sulfate, penicillin-streptomycin, EGF and FGF. U251 cells were produced and grown as previously described (Liu et al., 2019).

**Raman Spectroscopy.** Raman spectral concentration calibration was performed using bovine serum albumin, calf thymus DNA, S. cerevisiae RNA (Sigma Aldrich, St. Louis, MO, USA), and bovine heart lipids (Avanti Polar Lipids, Alabaster, AL, USA), with unit weight of 100 mg/mL for proteins and 20 mg/mL for RNA, DNA and lipids concentrations. The BCA toolbox yields a set of biomolecular weights (e.g., proteins, DNA, RNA, lipids and glycogen) and residual profile for each analyzed spectrum. Representative BCA processed spectra and output data are shown in Figure 1. The measurements were done as previously described.(Lita et al., 2019) Each data point is an average of at least 3 measurements.

**Organelle Extraction**. ER, Golgi and Lysosomes from approximatively 200 mg of wet pellet of $U251^{WT}$, $U251^{R132H}$ and $U251^{R132C}$ cells were extracted. Commercial kits described in the table were used for organelle extraction according to the manufacturer instructions (Invent Biotechnologies). Pellets were collected and stored in -80$^0$C for BCA protein quantification and subsequent metabolite extractions.

**LCMS global profile extraction.** 500 mL of chilled MilliQ water was added to each organelle pellet. A 40 µL aliquot was collected and stored at -80 $^0$C for BCA protein quantification. Organelle pellets were individually sonicated at 40 amps for 0.5 min. Bligh and Dyer biphasic liquid extraction was performed at 2:2:1.5 water/methanol/chloroform. Chilled 50 % MeOH (aq) reagent spiked with qualitative internal standards (qIS), nitrodracylic acid and isocaramidine sulfate was added to extract solution, vortexed and incubated on ice for 10 min. The chilled chloroform reagent spiked with qIS, phenyl-N-pyridinyl acrylamide was added to the extract solution and placed on rotating mixer for 60 min on ice. The samples were centrifuged at 12,000 rpm for 18 min at 4 $^0$C. The resulting two phases (upper hydrophilic and lower hydrophobic lipid) were separated while the remaining protein disk was discarded. Extracts were concentrated under N2 gas vapor, snap frozen with dry ice and stored at -80 $^0$C.

**Global lipidomics profiling using Ultra High-Performance Liquid Chromatography and Quadrupole Time-Of-Flight Mass Spectrometry (UHPLC-QTOF-MS)LC/MS lipidome acquisition.** The hydrophobic organelle extracts were reconstituted in reagent containing methanol: ACN: water as previously described (Altadill et al., 2017). Pooled quality control (QC) samples were composed of 10% volume from each sample. LC/MS lipidomic analysis was performed on acquired on the Agilent 6545 Quadrupole Time-of-Flight Mass Spectrometer coupled with Infinity II 1290 Liquid Chromatography Ultra-High-Pressure system. For each tissue extract using Acquity UPLC CSH 1.7 µm, 2.1 × 100 mm column (Waters Corp. Mass., USA, 186005297) using a gradient described by (Altadill et al., 2017).

**Enzyme prediction.** The differential enrichment analysis was performed via Metaboanalyst 4.0(Chong et al., 2018) platform for metabolites presenting fold-change ≥ 1.4. The hypergeometric p-values (p) were calculated based on probability of randomly selecting enzyme/pathway for over representation analysis (ORA) given the predicted metabolite set and the total number metabolites/substrates associated with the enzyme/pathway. Fold enrichment (FE)was determined based on hits from the predicted metabolite set divided by the hits expected by chance. The -log (p) and (FE) were performed to scale and plot data in Excel; where p is represented by color and FE is represented by size.

**LC/MS metabolome acquisition**. Metabolite extracts were resuspended in 60% methanol (aq) was acquired on the Agilent 6545 Qtof-MS with Infinity II 1290 UHPLC. LC/MS data acquisition was conducted with muliple polar assays developed to achieve broad detection and high resolution of amino acids, sugar phosphates and central carbon metabolites. Global profiling of polar metabolites and relative quantification for steady state, time-dependent $^{13}$C-label flux of polar metabolites was conduct on both, the AdvanceBio Glycan Map 2.1 x 150 mm 2.7 µm column (Agilent Technologies, Ca., USA, 683775-913) and InfinityLab Poroshell 120 HILIC-Z 2.1 x 100 mm, 2.7 µm column (Agilent Technologies, Ca., USA, 685775-924). Only LC/MS grade solvents and additives were used to prepare reagents, mobile phases and wash solutions, unless otherwise indicated. Wash cycles consisting of strong wash (50% Methanol, 25% Isopropanol, and 25% Water), weak wash (90% Acetonitrile and 1 % Water), and seal wash (10 % Isopropanol and 90 % water) were utilized to eliminate carryover for consecutive injections. Glycan Map column acquisition was performed in two experiments: both positive and negative electrospray ionization (ESI) modes. Compounds were resolved over a gradient composed of mobile phase A—10 mM ammonium acetate in 88% water and 12% acetonitrile, pH 6.85—and mobile phase B—10 mM ammonium acetate in 90 % acetonitrile(aq), pH 6.85—using a constant column temperature 25 $^{0}$C. The gradient was applied at flow rate 0.24 mL/min: 100 % B, hold 0.4 min; 97% B, min 1.0; 90 % B, min 3.5; 76 % B, min 6.5; hold 0.25 min; 75 % B, min 7.65; 74.8 % B, min 8.25; hold 0.5 min; 72 % B, min 9.25; 72.1 % B min 9.75, 97 % B min 10.5; re-equilibrate for 2 min. HILIC-Z column acquisition was performed ESI negative mode. Compounds were resolved over a gradient composed of mobile phase A—10 mM ammonium acetate in 88% water and 12% acetonitrile, pH 6.85—and mobile phase B—10 mM ammonium acetate in 90 % acetonitrile (aq)—using a constant column temperature 30 $^{0}$C. The gradient was applied at flow rate 0.24 mL/min: 100 % B, 0.5 min; 95% B, 2.0 min; 60 % B, 3.0 min; 35 % B, 5 min; hold 0.25 min; 0% B, 6 min; hold 0.5 min; 100 % B, 7.25 min. The mass analyzer parameters included drying gas temperature 250 $^{0}$C, sheath gas temperature 325 $^{0}$C, nebulizer 45 psi, skimmer 50 V, octopole radio frequency 750 V and scan rate of 4 spectra/s. In ESI positive mode experiment, ms spectra were acquired over a voltage gradient of capillary 3500 V, nozzle 2000 V, and fragmentor 165 V. In ESI negative mode experiment, mass spectra were acquired over a voltage gradient of capillary 3000 V, nozzle 1500 V, and fragmentor 80 V.

**LC/MS data analysis:** Prior to preprocessing each dataset, pooled QC samples (TIC, BPI and EIC) were chromatographically examined to inspect consistency of retention time and ionization levels throughout. Following acquisition, mass feature bins were defined by partitioning the m/z vs. retention time (RT) matrices into fixed width using Agilent Masshunter Profinder B.08.00. Bins were manually inspected to confirm consistent, reproducible integration for each compound of interest across all samples. Precursor m/z for each bin was determine using molecular feature extraction algorithm to deconvolute, integrate, and envelope parent ions, adducts (H-, Cl+, H+, Na+), natural isotopes and neutral losses to define each composite spectrum. Logical binning of the input mass data was conducted via targeted ion selection, annotation and alignment restricted to accurate neutral mass ±5.0 mDa with retention time ±0.4 min of references defined in Personal Compound Data Library (PCDL). Following pre-processing, the ion abundance for each sample were corrected using sample-specific mass quantification. Values were corrected to sample mass and internal standard response. Comparative analysis and graphs were generated in Excel Office 16. For each graph, features include standard deviation between 3 technical replicates from each sample and represent mean comparison as relative %. In order to fit relative % graph and dispersion of heterogenous classes of fatty acids, the mean abundance of each feature was scaled based on percent of feature with greatest mean abundance in each set.

**Protein Digestion and TMT labeling:** The fractionated subcellular fractionated pellets were solubilized in 50 mM HEPES, pH 8.0 containing 20% methanol. Digestion was performed by addition of trypsin at a ratio of 1:50 (Promega) and incubating overnight at 37$^{o}$C. Digestion was acidified by adding formic acid (FA) to a final concentration of 1%. The digest was desalted using Pierce peptide desalting columns according to manufacturer's protocol. Peptides were eluted from the columns using 50% ACN/0.1% FA, dried in a speedvac and kept frozen in -20$^{o}$C for further analysis. The concentration of the peptide was estimated using Pierce Quantitative Fluorescent peptide assay kit.

For TMT labeling 25ug of each sample was reconstituted in 50 ul of 50mM HEPES, pH 8.0, and 100ug of TMT label in 100% ACN was added to each sample. After incubating the mixture for 1 hr at room temperature with occasional mixing, the reaction was terminated by adding 8 ul of 5% hydroxylamine. The peptide samples for each subcellular fraction were pooled and speedvac to dry labeled peptide sample. The samples were cleaned up using peptide desalting columns.

**High pH reverse phase fractionation:** The first dimensional separation of the peptides was performed using a Waters Acquity UPLC system coupled with a fluorescence detector (Waters, Milford, MA) using a 150mm x 3.0mm Xbridge Peptide BEM$^{TM}$ 2. 5 um C18 column (Waters, MA) operating at 0.35 ml/min. The dried peptides were reconstituted in 100 ul of mobile phase A solvent (3 mM ammonium bicarbonate, pH 8.0). Mobile phase B was 100% acetonitrile (Thermo Fisher). The column was washed with mobile phase A for 10 min followed by gradient elution 0- 50% B (10-60 min) and 50-75 %B (60-70 min). The fractions were collected every minute. The fractions collected along the fractionation were pooled into 24 fractions, vacuum centrifuged to dry and stored at -80$^{o}$C until analysis by mass spectrometry.

**Mass Spectrometry acquisition and data analysis:** The dried peptide fractions were reconstituted in 0.1%TFA and subjected to nanoflow liquid chromatography (Thermo Easy nLC 1000, Thermo Scientific) coupled to high resolution tandem MS (Q Exactive, HF, Thermo Scientific). Peptides were separated using a second dimension low pH gradient using a 2-40% ACN over 120 minutes in mobile phase containing 0.1% formic acid at 300 nl/min flow rate. MS scans were performed in the Orbitrap analyser at a resolution of 120,000 with an ion accumulation target set at 3e$^{6}$ and max IT set at 50ms over a mass range of 200-1800 m/z, followed by MS/MS analysis at a resolution of 45,000 with an ion accumulation target set at 1e$^{5}$, max IT of 120ms and first fixed mass set at 105 m/z. MS2 precursor isolation width was setup at 0.7 m/z, normalized collison energy was 29, and charge state 1 and unassigned charge states were excluded.

Acquired MS/MS spectra were searched against a human uniprot protein database along with a contaminant protein database, using a SEQUEST and percolator validator algorithms in the Proteome Discoverer 2.2 software (Thermo Scientific, CA). The precursor ion tolerance was set at 10 ppm and the fragment ions tolerance was set at 0.02 Da along with methionine oxidation included as dynamic modification and TMT6 plex (229.163Da) set as a static modification of lysine and the N-termini of the peptide. Trypsin was specified as the proteolytic enzyme, with up to 2 missed cleavage sites allowed. Searches used a reverse sequence decoy strategy to control for the false peptide discovery and identifications were validated using percolator software.

**Confocal Microscopy:** A Zeiss LSM880 confocal microscope equipped with a 63x plan-apochromat (N.A. 1.4) oil immersion objective lens, a PeCon stage top incubator to control temperature (37C), humidity and CO2, and a Bioptechs objective lens heater was used to acquire confocal images of live U251 IDH1 mutant and $IDH^{WT}$ cells labeled with RFP expressing proteins in different organelles and C-16 BODIPY for saturated fatty acid co-localization. Confocal images were collected with 2x frame averaging, 1.0 um optical section thickness and 0.09 um X-Y pixel size. A corresponding differential interference contrast (DIC) image was also acquired. A Nikon Ti2-E microscope equipped with a Yokogawa CSU-W1 spinning disk confocal unit, a 60x plan-apochromat (N.A. 1.4) oil immersion objective lens and Hamamatsu ORCA Flash 4.0 V3 sCMOS camera was used to acquire images of live $U251^{WT}$, $U251^{R132H}$, $U251^{R132C}$ cells labeled with lysotracker and in absence and presence of $IDH1^{mut}$ inhibitor, AGI-5198. Time-lapse confocal images were collected every second over a 120 second time period with 0.1 um X-Y pixel size. Lysosome displacement was quantified using the spot tracking module of Imaris (v.9.2) image processing and analysis software. A Zeiss Elyra 7 lattice structured illumination microscope equipped with a 63x alpha plan-apochromat (N.A. 1.46) oil immersion objective lens and dual PCO Edge 4.2 sCMOS cameras was used to acquire z-stacks of $U251^{R132H}$ cells labeled with Golgi RFP and C16-BODIPY. Images were collected using fast frame switching in lattice SIM mode with 0.1 um z-step size, 0.03 um X-Y pixel size, and processed using the SIM module of the Zen software.

**Transmission Electron Microscopy (TEM)**:The $U251^{WT}$, $U251^{R132C}$ and $U251^{R132H}$ cell lines were cultured in 6 well plates and fixed in glutaraldehyde (2% v/v) cacodylate buffer (0.1M, pH 7.4) for at least 2 hrs in preparation for thin-sectioned EM analysis previously described (Nagashima et al., 2011). Patient tissue was stored fresh in formaldehyde (4% v/v), glutaraldehyde (2% v/v) cacodylate buffer (0.1M, pH 7.4) for at least 2 hrs. Processing and embedding were carried out at room temperature in a fume hood. The cells were washed 2 times in cacodylate buffer prior to post-fixation of 1hr in osmium tetroxide (1% v/v). *en bloc* stain in 0.5% w/v uranyl acetate (0.5% v/v) in acetate buffer (0.1M, pH 4.5) for 1hr and in the dark to prevent the uranyl acetate from being precipitated. The cells were dehydrated through a series of multiple washes of ethanol solution (35%, 50%, 75%, 95%, and 100%). Following 3 washes of 100% ethanol, the cells were washed with pure epoxy resin overnight and washed 2 more times the following day prior to embedding. Once embedded the resin was cured for 48 hrs at 55$^{o}$C. The cured resin blocks were separated from the plate by submerging in liquid nitrogen. In preparation for thin sectioning, the separated resin blocks were examined under an inverted microscope to select an area with a large number of cells. The selected area was thin sectioned at 100nm and mounted on 150 copper mesh grids. The grids were counter stained in uranyl acetate and lead citrate and carbon coated in a vacuum evaporator. The grids were scanned and imaged at high and low magnification in the electron microscope operated at 80kv. A CCD camera captured the digital images.

**EC50 determinations:** Sensitivity of patient-derived glioma cell lines to oleic acid was detected using the cell viability, CCK-8 kit. Various types of patient-derived glioma cell line were seeded as single cell suspension at a concentration of 20,000 cells/well in 96-well plates. Cells were treated with oleic acid (Sigma Cat#75090) at concentrations ranging from 20 µM to 400 µM or DMSO as control and incubated for 72 hours. Relative cell numbers were quantified by the CCK-8 assay (Dojindo Molecular Technologies, Rockville, MD) according to the manufacturer's protocol. All assays were performed in five replicates.

**Apoptosis assay using flow cytometry:** Apoptosis was assessed using PE Annexin V Apoptosis Detection Kit I (BD Biosciences) and analyzed by flow cytometry. Briefly, cells were plated into 6-well culture dishes ($1\times10^{6}$ cells/well) for 24 hrs. prior to the addition of oleic acid (Sigma Cat#75090) at a concentration of 150 µM. Following 24 hrs incubation with oleic acid the percentage of apoptotic cells was determined by the annexin V-PE/7-AAD assay following manufacturer's instructions. Fluorescence of the cells was immediately determined by a Sony SA3800 spectral analyzer.

**SiRNA.** $5\times10^{5}$ cells/well in 6-well plates were transfected with a set of three siSCD1 specific-targeting small interfering RNAs (OriGENE Inc, SR304248) using siTran 2.0 siRNA transfection reagent (OriGENE Inc, TT320001) for 18h following the manufacturer's protocol. Trilencer-27 Fluorescent-labeled transfection control siRNA duplex (OriGENE Inc, SR30002) were used as controls. After three days incubation, cells were treated again with specific-targeting siRNA or control siRNA for 2 days and then analyzed.

**RT-PCR.** Total RNA was extracted by using RNeasy spin columns (Qiagen, Chatsworth, CA, USA; cat. no.: 74136). After treatment with RNase-free DNase (Qiagen), RT was performed on RNA by using Clontech RNA to cDNA EcoDry kit (cat.no.: 639548) according to the manufacturer's protocol. RT-qPCR was performed on QuantStudio. Reactions were carried out in 10 $\mu$L by using SYBR Green PCR master mix according to the manufacturer's protocol (Bio-Rad, cat. no.: 1725017, Hercules, CA). The concentration of primer pairs was 100 nM. All primer sequences are shown in the primer table. All reactions were performed in triplicate. To make comparisons between samples and controls, the CT (cycle threshold, defined as the cycle number at which the fluorescence is above the fixed threshold) values were normalized to the CT of *beta*-actin in each sample. The following primers were used hSCD1-F: 5'-AAACCTGGCTTGCTGATG-3'; hSCD1-R: 5'-GGGGGCTAATGTTCTTGTCA-3'; beta-actin-F: 5'-ACTGGAACGGTGAAGGTGAC-3', beta-actin-R: 5'-GTGGACTTGGGAGAGGACTG-3'

### SUPPLEMENTARY FIGURE LEGENDS

**Supplementary Figure S1: Detailed analyses of organelle specific Raman data shows that the composition of lipids is different in different mutants. a.** Different mutants produce different concentration of D-2hydroxyglutarate as measured via LC/MS. **b.** Comparison of average levels of biomolecular components obtained from Raman analysis show differences due to the R132H, and R132C mutations. The levels above the x axis show increased levels in the U251$^{R132H/C}$ cells, while the bars below the x axis show increased levels in the U251$^{WT}$ cells. C denotes U251$^{R132C}$, while H denotes U251$^{R132H}$ cells. **c** and **d.** Raman spectrum depicting the assignment from which information regarding lipid structure is obtained such as the lipid unsaturation parameter and the Cis/Trans ratio and various Raman spectra for standards used. **e-g.** Sphingomyelin levels are heterogeneously distributed in organelles of both mutant cells (upper panels) and become more homogenous after addition of AGI-5198 inhibitor with the exception of lysosomes (lower panels).

**Supplementary Figure S2: Heterogenous distribution of lipid parameters across cells and organelles. a-c.** Cholesterol levels are heterogeneously distributed in both U251$^{R132H}$ and U251$^{R132C}$ cells compared with the U251$^{WT}$ cells (upper panels) and this distribution become more homogenous upon addition of AGI-5198 inhibitor (lower panels) with the exception of lysosomes. **d-f** LSU parameter is heterogeneously distributed in both U251$^{R132H}$ and U251$^{R132C}$ cells compared with the U251$^{WT}$ cells (upper panels) and this distribution become more homogenous upon addition of AGI-5198 inhibitor (lower panels) with the exception of lysosomes. **g-i.** TCP parameter is heterogeneously distributed in both U251$^{R132H}$ and U251$^{R132C}$ cells compared with the U251$^{WT}$ cells (upper panels) and this distribution become more homogenous upon addition of AGI-5198 inhibitor (lower panels) with the exception of lysosomes.

**Supplementary Figure 3.** Lysosomes increase in number, accumulate lipids and traffic faster in presence of AGI-5198 inhibitor. **a**. Confocal microscopy shows increased number of lysosomes in U251 $^{R132H}$ (middle panel) and U251$^{R132C}$ (lower panel). **b**. Confocal microscopy shows significant increase in lysosomes (labeled with lysotracker, red) in presence of 12 uM of AGI-5198 in all samples. **c** and **d.** Transmission electron micrographs displayed increaded lysosomal content in U251$^{WT}$ (upper panel), U251$^{R132C}$ (middle panel) and U251$^{R132H}$(lower panel) upon addition of 12 mM AGI-5198. **e-g**. Samples from oligodendroglioma patient tissue (**e** and **f**) displayed increased lysosomal content compared with the content of lysosomes in glioblastoma multiforme tissue (**g**). **h**. Western Blot analysis of LAMP-1 confirms the increased lysosomes in mutant cells compared with U251$^{WT}$ and increased lysosome in all samples upon addition of AGI-5198. **i**. Lysosome area was significantly increased only in U251$^{R132H}$ when compared with U251$^{WT}$ cells. j. Raman-based measurements show increase in total lipids in lysosomes of U251$^{R132H}$ and U251 $^{R132C}$ when compared with U251$^{WT}$ cells. Addition of AGI-5198 increased lysosomal lipids further. K. representative image of lysosomal movement tracking in U251 R132H cells compared with U251WT cells. **l** and **m**. Quantification of lysosomal speed via mean average displacement. All lysosomes were considered for these measurements, although there is a clear population of lysosomes moving fast. (supplemental Movie 1).

**Supplementary Figure S4**. **Confocal microscopy reveals no co-localization between SFA and ER, mitochondria or lysosomes within the timeframe of the experiment**. Confocal Microscopy showing no signs of co-localization of SFAs (BODIPY™ FL C16, green) at ER membrane (labeled with RFP-BacMan 2) (**a**) mitochondria (**b**) (labeled with RFP BacMan 2) or lysosomes (**c**) (labeled with RFP BacMan 2).

**Supplementary Figure S5. SCD-1 and SCD-5 overexpression is not correlated with survival of in astrocytoma or glioblastomas irrespective of the presence of IDH1 mutation. a-d)** Kaplan-Meier survival plots of patients with astrocytoma and GBM and different expressions of SCD-1. SCD-1$^{high}$ is depicted in orange while SCD-1$^{low}$ in cyan. **e-h)** Kaplan-Meier survival plots of patients with astrocytoma and GBM and different expressions of SCD-5. SCD-5$^{high}$ is depicted in orange while SCD-5$^{low}$ in cyan.

**Supplementary Figure S6**. **SCD-1 as the major player in lipid imbalance and MUFA-induced cell death. a**. Western Blot analysis of fatty acid synthesis and regulation by SREPB1 showed increased SCD-1 protein levels in IDH1$^{mut}$ patient-derived cell lines, BT142, TS603 and NCH1681 and low expression of this protein in GSC827 and GSC923 (IDH1$^{WT}$). **b**. mRNA levels of SCD-1 relative to $\beta$-actin of GSC827 (IDH1$^{WT}$) and TS603 (IDH1$^{mut}$ oligodendroglioma). **c**. Addition of oleic acid leads to visible cell death as shown in the images of TS603 (upper panels), BT142 (middle panels) and NCH1681 (lower panel). **d**. Oleic acid addition does not lead to apoptosis in GSC923 cells. **e**. Inhibition of Fatty acid synthase (FASN) leads to more specific growth inhibition on U251$^{WT}$ cells. **f**. Addition of linoleic acid (PUFA) affects the growth of U251$^{WT}$ cells specifically. **g**. Oil red stain confirms the accumulation of neutral lipids in the lipid droplets formed by the oleic acid addition.

**Supplementary Figure S7**. The effect of deferoxamine on a) IDH1$^{mut}$ cells: TS603 (upper graph), BT142 (lower graph) and b) IDH1$^{WT}$ cells: GSC827 (upper graph) and GSC923 (lower graph).

### SUPPLEMENTARY FIGURES

**Supplementary Figure S1:**

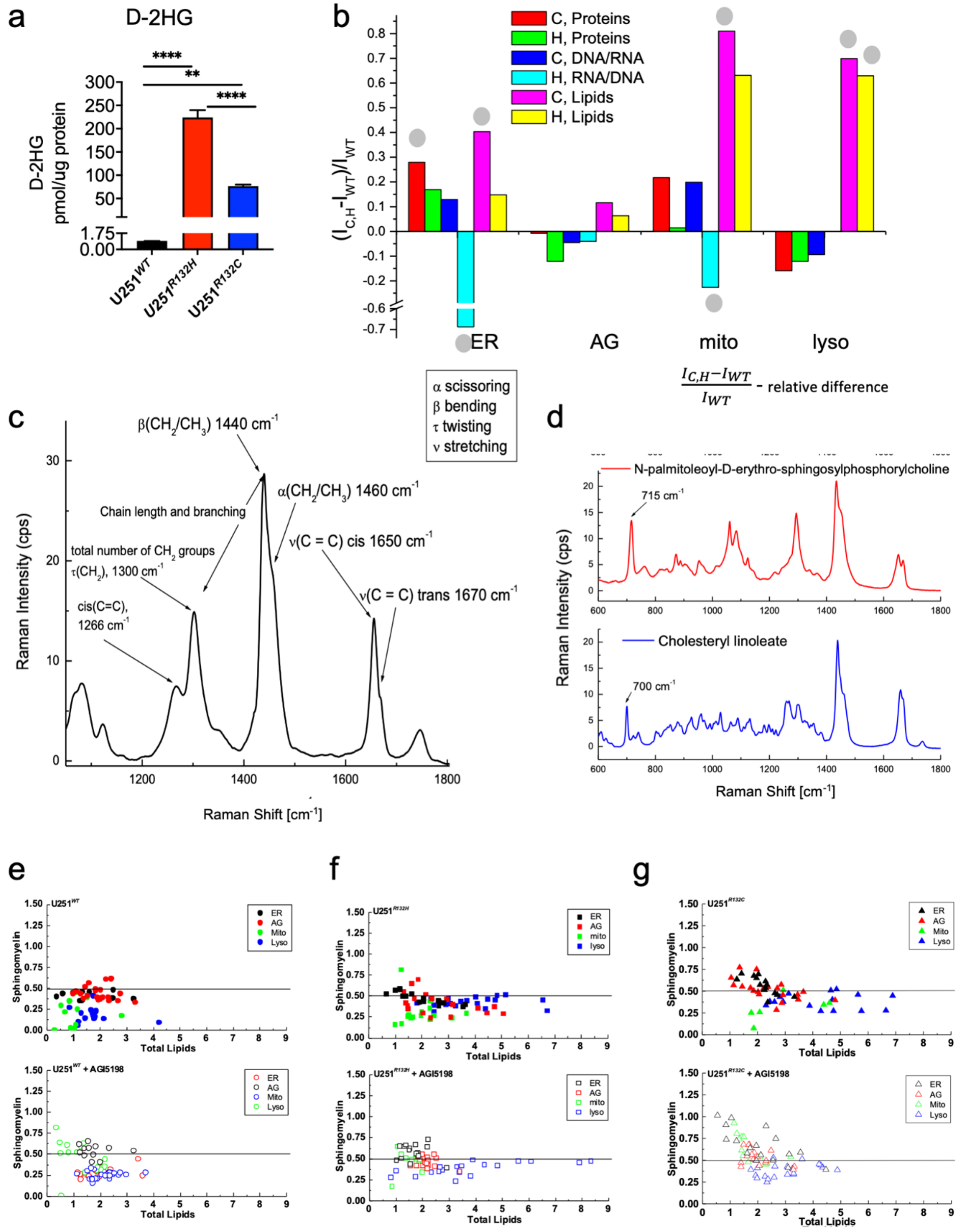

**Supplementary Figure S2:**

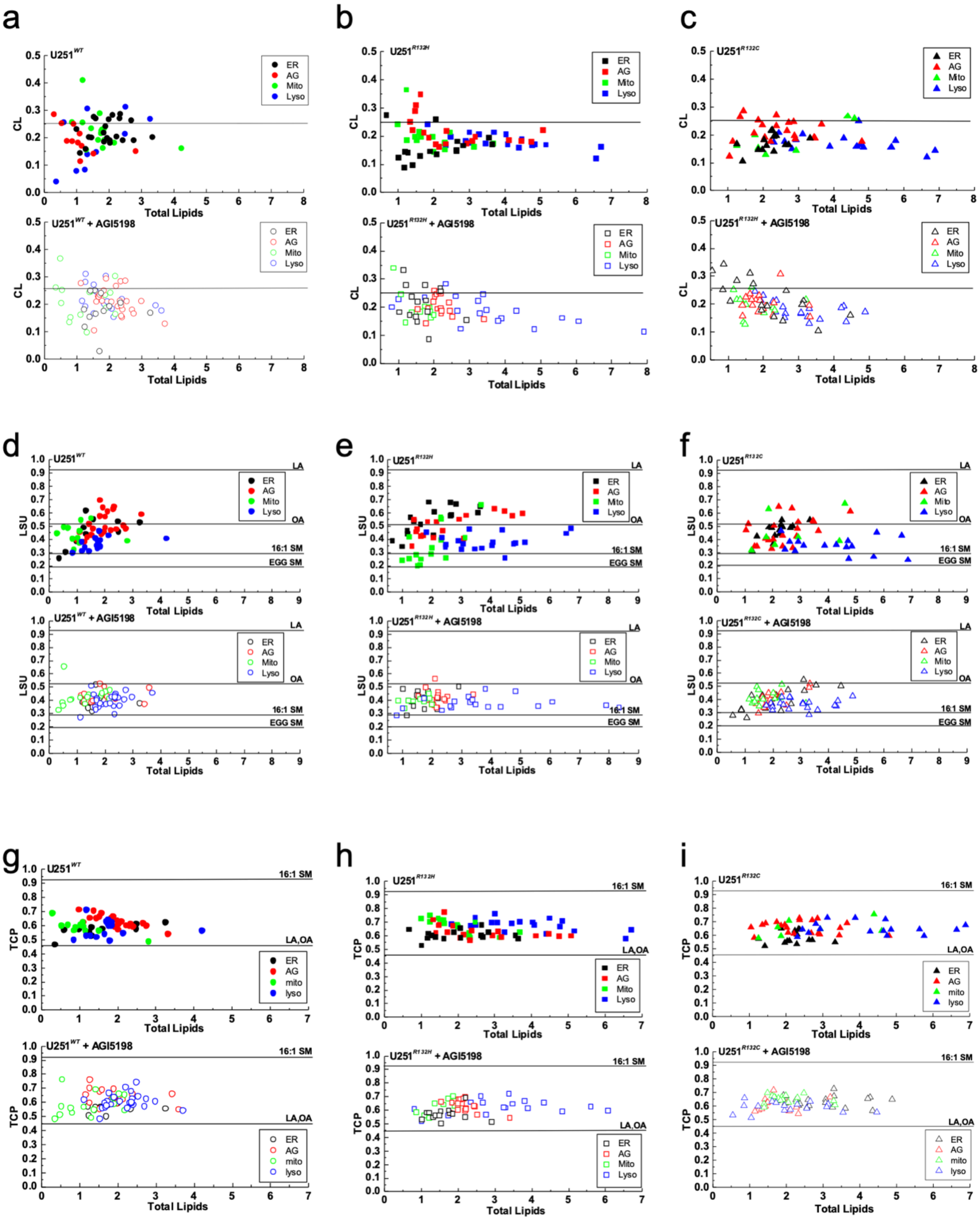

**Supplementary Figure S3:**

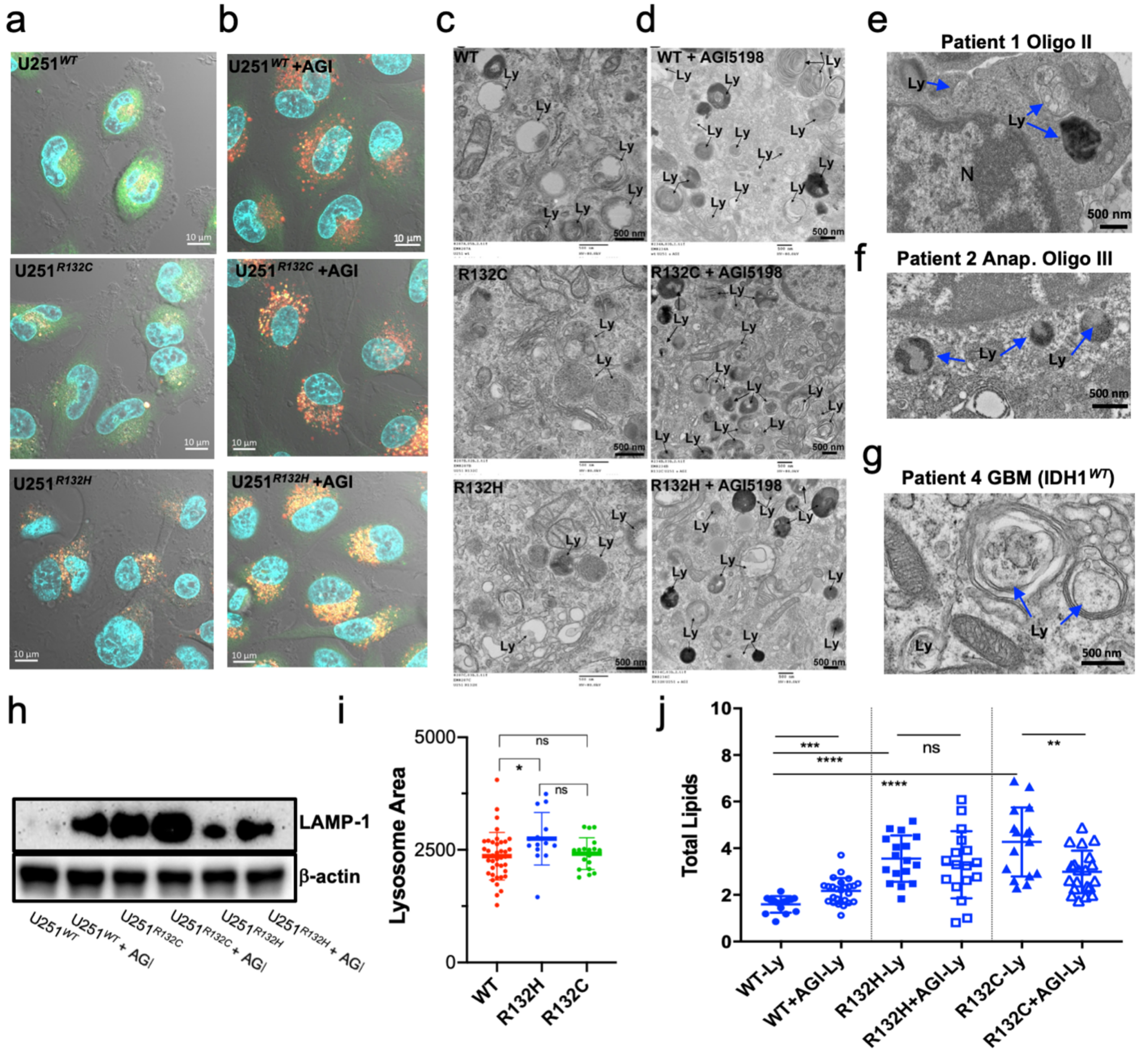

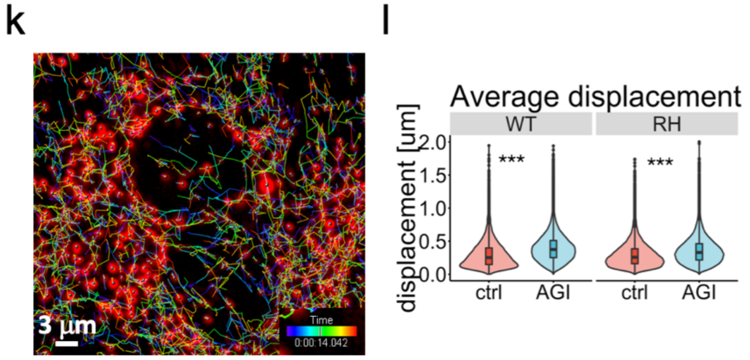

m

| Sample | Mean Displacement (um/s) |
|---|---|
| U251WT | 0.3 |
| U251R132H | 0.3 |
| U251WT+AGI | 0.4 |
| U251R132H+AGI | 0.36 |

**Supplementary Figure S4:**

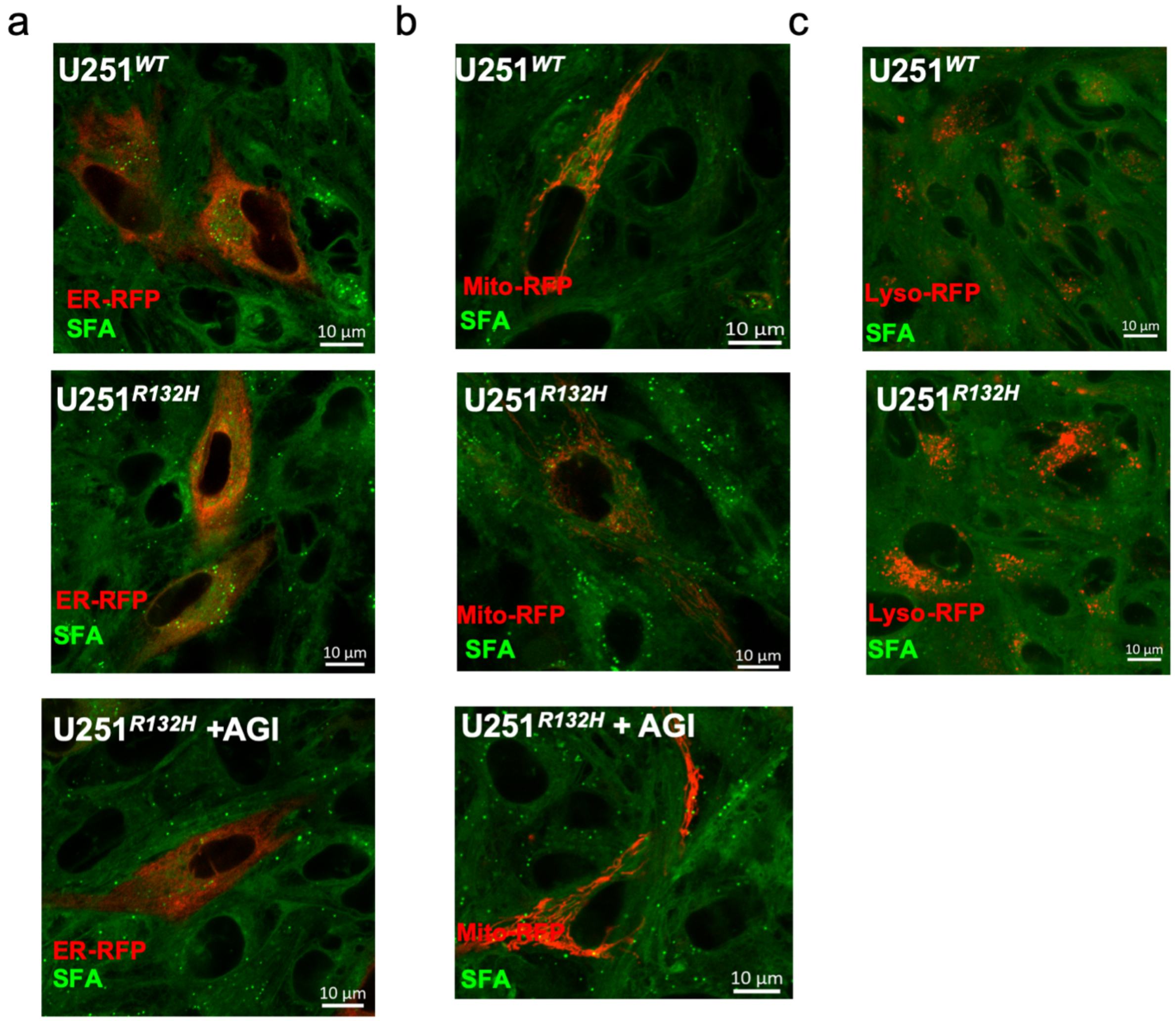

**Supplementary Figure S5:**

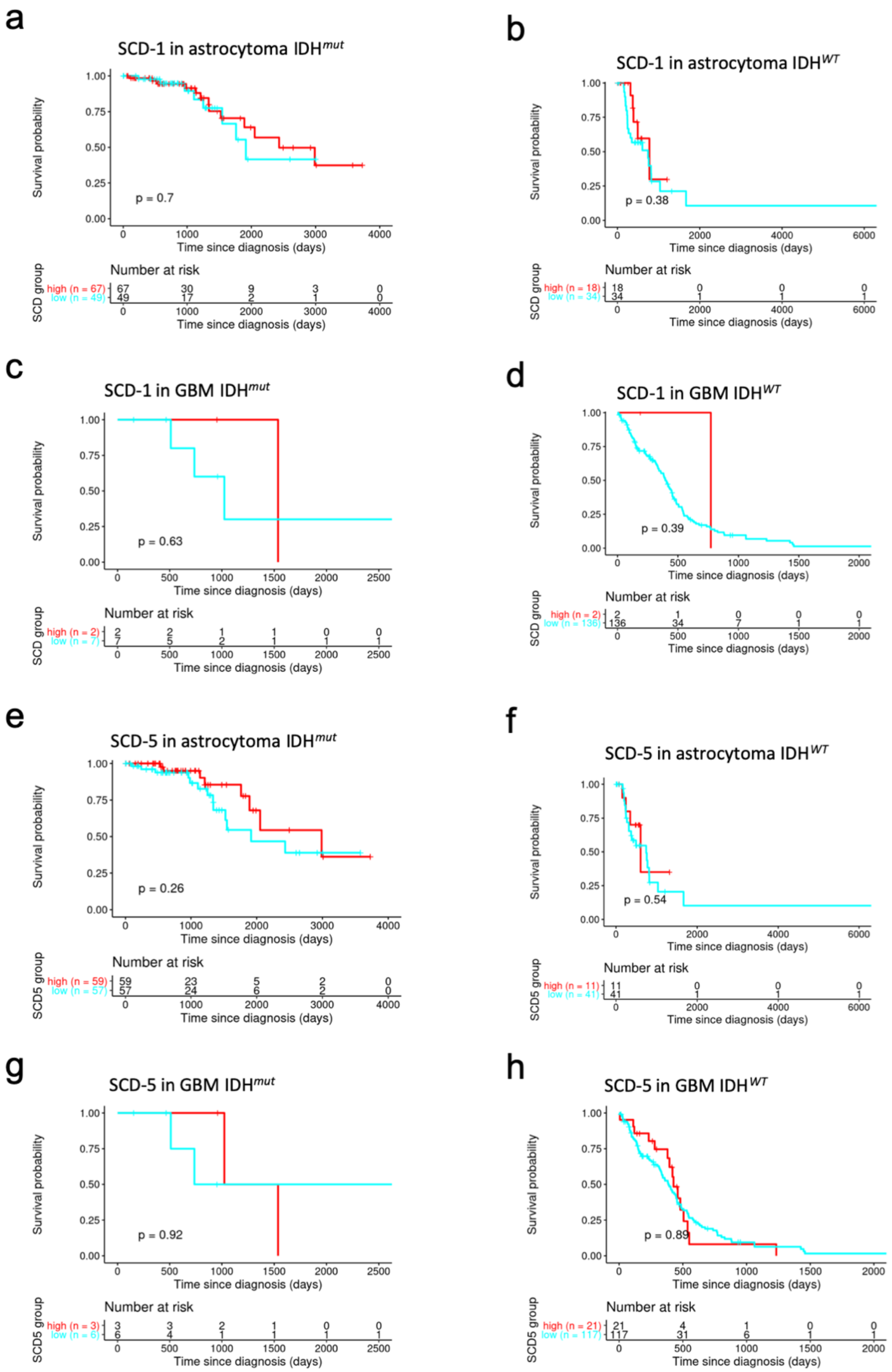
a
SCD-1 in astrocytoma IDH$^{mut}$
p = 0.7
b
SCD-1 in astrocytoma IDH$^{WT}$
p = 0.38
c
SCD-1 in GBM IDH$^{mut}$
p = 0.63
d
SCD-1 in GBM IDH$^{WT}$
p = 0.39
e
SCD-5 in astrocytoma IDH$^{mut}$
p = 0.26
f
SCD-5 in astrocytoma IDH$^{WT}$
p = 0.54
g
SCD-5 in GBM IDH$^{mut}$
p = 0.92
h
SCD-5 in GBM IDH$^{WT}$
p = 0.89
Survival probability
Time since diagnosis (days)
Number at risk
SCD group
SCD5 group

**Supplementary Figure S6:**

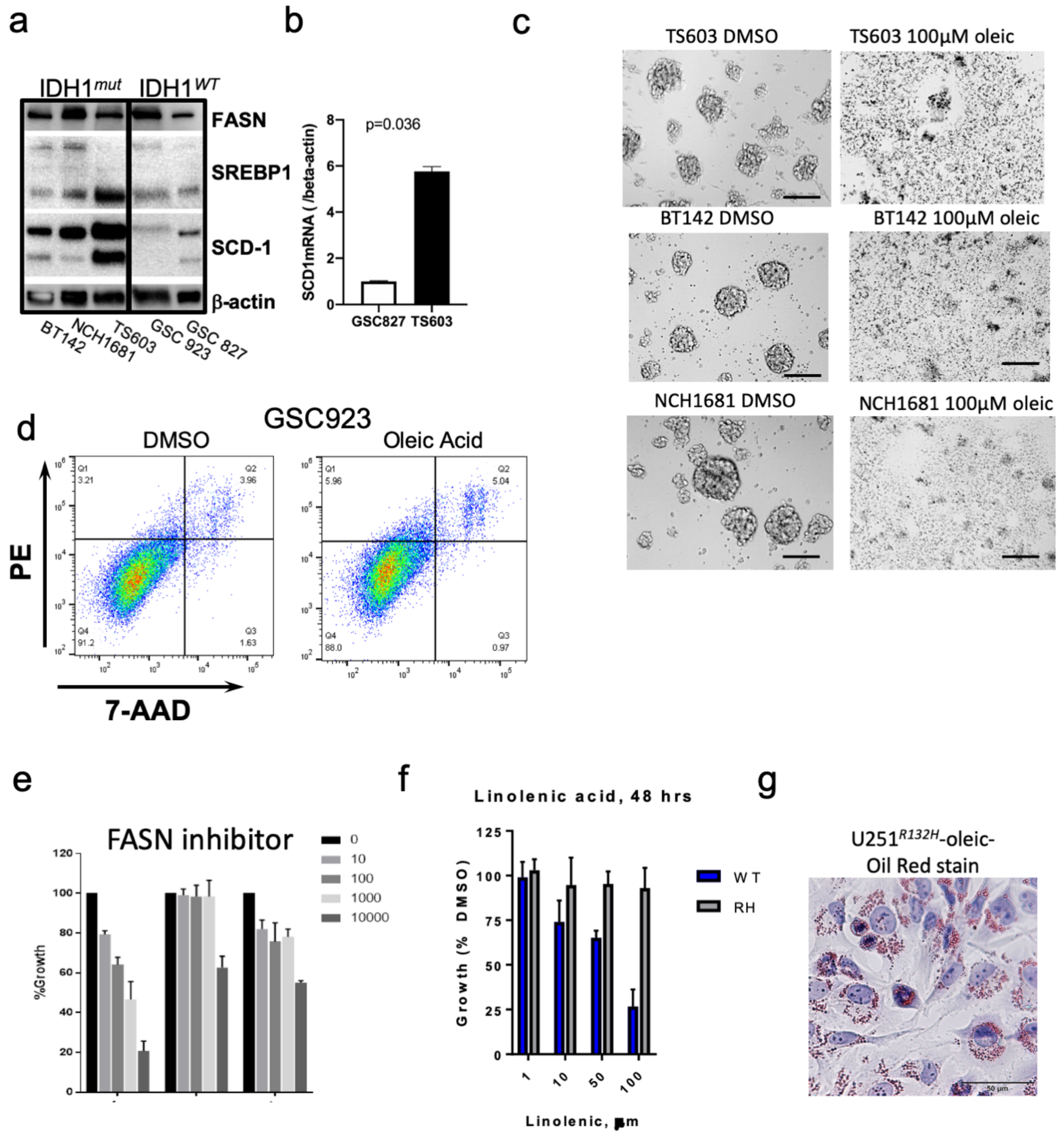

**Supplementary Figure S7:**

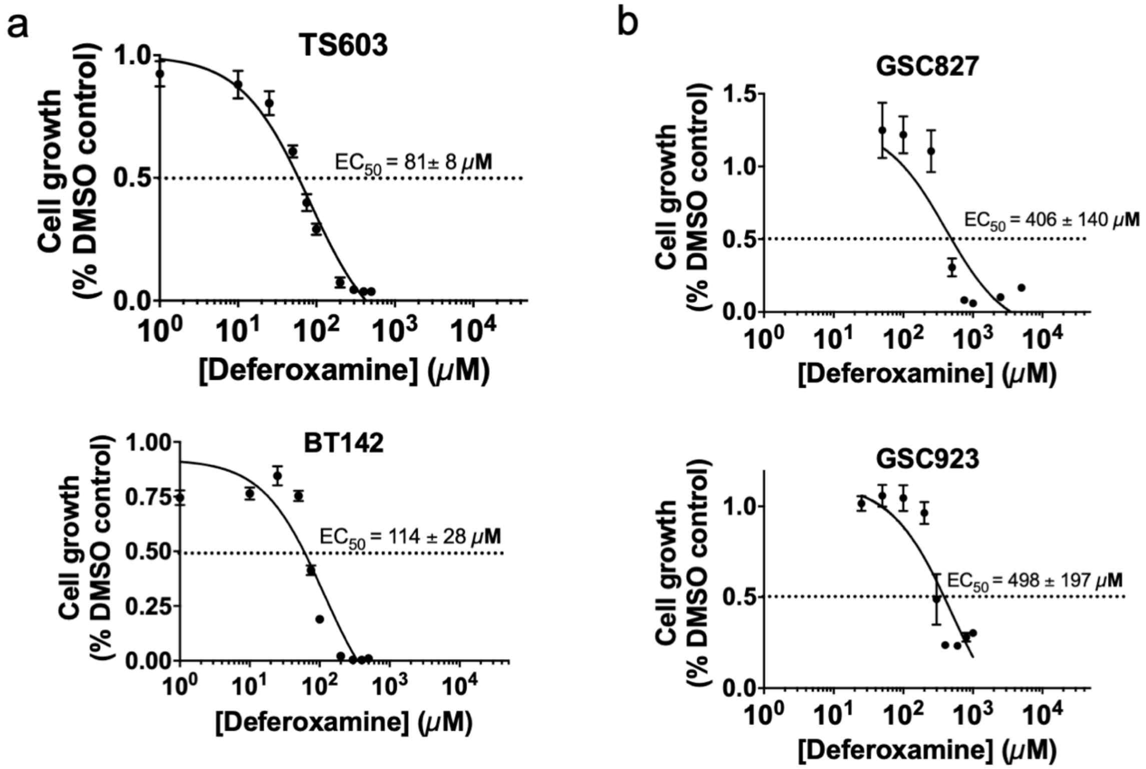
